# Sensory Entrained TMS (seTMS) enhances motor cortex plasticity

**DOI:** 10.1101/2025.07.23.666433

**Authors:** J.M. Ross, L. Forman, U. Hassan, J. Gogulski, J. Truong, C.C. Cline, S. Parmigiani, N-F. Chen, J.W. Hartford, T. Fujioka, S. Makeig, A. Pascual-Leone, C.J. Keller

**Affiliations:** Department of Psychiatry and Behavioral Sciences, Stanford University Medical Center, 401 Quarry Road, Stanford, CA, 94305, USA; Wu Tsai Neurosciences Institute, Stanford University, Stanford, CA, USA; Veterans Affairs Palo Alto Healthcare System, and the Sierra Pacific Mental Illness, Research, Education, and Clinical Center (MIRECC), 3801 Miranda Avenue, Palo Alto, CA 94304, USA; Department of Clinical Neurophysiology, HUS Diagnostic Center, Clinical Neurosciences, Helsinki University Hospital and University of Helsinki, Helsinki, FI-00029 HUS, Finland; Department of Neuroscience and Biomedical Engineering, Aalto University School of Science, Rakentajanaukio 2, 02150, Espoo, Finland; Center for Computer Research in Music and Acoustics (CCRMA), Department of Music, Stanford University, Stanford, CA, USA; Swartz Center for Computational Neuroscience, Institute for Neural Computation, University of California, San Diego, CA, USA; Department of Neurology, Harvard Medical School, Boston, MA, USA; Deanna and Sidney Wolk Center for Memory Health, Hebrew Senior Life, Hinda and Arthur Marcus Institute for Aging Research, Boston, MA, USA

**Author notes:** **Correspondence and requests for materials should be addressed to JMR.** Jessica M. Ross, PhD, Stanford University, Department of Psychiatry and Behavioral Sciences, 401 Quarry Road, Stanford, CA 94305-5797.

**Keywords:** Non-invasive brain stimulation (NIBS), Transcranial magnetic stimulation (TMS), Electroencephalogram (EEG), Motor Evoked Potential (MEP)

## Abstract

Neural excitability fluctuates with sensory events, creating windows of opportunity to enhance brain stimulation. Repetitive transcranial magnetic stimulation (TMS), including intermittent theta burst stimulation (iTBS), is a promising treatment for neurological and psychiatric disorders, but does not account for fluctuations in neural excitability, likely contributing to variable outcomes. Sensory Entrained TMS (seTMS) leverages sensorimotor oscillations to enhance corticospinal responses, but the sustained effects as a repetitive protocol are unknown. We extended seTMS to iTBS measuring motor-evoked potentials (MEPs) as a physiological readout in a randomized crossover study comparing standard iTBS with sensory entrained iTBS (se-iTBS, *n*=20). se-iTBS more than doubled the MEP effect (55% vs. 26% MEP enhancement) and persisted for at least 30 minutes. Notably, more than 80% of participants showed larger responses with se-iTBS at all time points. se-iTBS may provide a robust and practical framework for optimizing TMS that bridges electrophysiological mechanisms and clinical applications.

## 1. Introduction

Non-invasive brain stimulation provides a unique opportunity to probe and alter neural circuits, yet the field faces substantial challenges due to highly variable stimulation effects. Transcranial magnetic stimulation (TMS) is a widely used form of noninvasive brain stimulation that delivers electromagnetic pulses to the scalp to induce currents in underlying cortical networks^1^. Repetitive TMS protocols, including intermittent theta burst stimulation (iTBS), have been shown to increase motor evoked potential (MEP) size^2^. This increase in MEP size is thought to reflect corticospinal plasticity, supported by evidence in rat models^3^ and non-human primates^4^. iTBS is used clinically for stroke recovery and various other treatments^5–8^. In stroke recovery, iTBS has demonstrated benefits for increasing motor excitability^9,10^, recovering motor function^11,12^, increasing functional connectivity in language networks^13^, increasing motor plasticity^12^, and improving daily functioning^11,12^ (see ^12^ for a systematic review and meta-analysis). iTBS to motor cortex has also shown promise for treating chronic pain, essential tremor, and dystonia^14^. Beyond motor applications, prefrontal repetitive TMS has demonstrated efficacy for medication-resistant depression in sham-controlled trials^15^ (though not yet against an active control site), and iTBS has achieved FDA-clearance as a shorter duration alternative7,^16,17^, with accumulating evidence supporting the antidepressant effects^17,18^, and with both thought to work by inducing changes in cortical excitability in prefrontal networks and promoting neural plasticity^19–21^.

Despite widespread clinical adoption of iTBS, substantial variability in neural responses^22–24^ and treatment outcomes^5,16^ remains a critical challenge. More insights from basic research must be used to improve the treatment protocols. For instance, baseline corticospinal excitability and electroencephalographic (EEG) oscillations significantly impact plasticity effects of iTBS^24–26^, but these findings have not been leveraged into better clinical protocols. The clinical impact would be substantial: traditional daily TMS depression treatments achieve remission in fewer than half of patients^5^, and newer accelerated treatments achieve remission one month after treatment in fewer than half of patients^16^. These rates underscore the pressing need for more reliable and effective protocols. This challenge has motivated more basic research on enhancing the effects of TMS. Given that EEG oscillations interact with TMS responses, leveraging these neural rhythms represents a promising avenue for enhancing TMS effectiveness.

Brain state dependent neuromodulation is a method that aligns TMS delivery with endogenous neural oscillations^25–28^. These methods use EEG to trigger TMS pulses during optimal phases of sensorimotor rhythms (mu, µ), demonstrably enhancing brain responses and plasticity as measured by MEPs^25,29,30^. However, clinical adoption remains limited due to the additional costs and complexity of real-time EEG integration. Therefore, a method for brain state dependent stimulation that circumvents EEG requirements would democratize these improved techniques for widespread clinical use.

Music offers a promising solution because passively listening to musical rhythms induces reliable changes in EEG oscillations^31–36^. EEG oscillatory phase changes are time-locked to musical events that are rhythmically predictable^33,37^, occur in multiple frequency bands^31–33,38–40^ across many brain regions^39^, and have been shown to reflect dynamically shifting excitability brain states^33,34,37,38,40^. Importantly, music that induces more sensorimotor coupling can be used to evoke larger TMS responses^41^. Clinical research has demonstrated that music-based interventions can be safely and effectively implemented in therapeutic settings^42,43^.

Based on these principles, we developed Sensory Entrained TMS (seTMS), which pairs music with TMS to synchronize relevant brain oscillations without requiring EEG. This approach strategically delivers TMS pulses when inhibitory oscillations are naturally desynchronized, and our prior study showed that this enhanced neural excitability as measured by larger MEPs^44^. Here we extend this work to iTBS by developing Sensory Entrained iTBS (se-iTBS), which pairs music with iTBS to leverage auditory entrainment for neural-synchronized brain stimulation. We hypothesized that se-iTBS would enhance plasticity effects compared with standard iTBS by optimally timing stimulation with neural excitability. If successful, this approach would offer a practical and accessible method for state dependent neuromodulation, with broad implications for therapeutic TMS for neurological and psychiatric disorders.

## 2. Methods

### 2.1. Participants and Study Design

This study was reviewed and approved by the Stanford University Institutional Review Board, performed in accordance with all relevant guidelines^45^ and regulations (including the Declaration of Helsinki), and written informed consent was obtained from all participants. 22 healthy participants (22-64 years old [M=42.6, SD=14.7, 12F/10M]) responded to an online recruitment ad and, after an initial online screening and consent, 20 of those were eligible to participate (22-64 years old [M=41.0, SD=14.5, 10F/10M, handedness 19R/1L]) and were enrolled. See Table S1 for more demographics. Of the 2 who were not enrolled, 1 was excluded due to scheduling conflicts and 1 due to the exclusionary criteria.

Inclusion criteria on the online screening form were (a) aged 18-65, (b) able to travel to study site, (c) fluent in English and (d) fully vaccinated against COVID-19. Exclusion criteria were (a) lifetime history of psychiatric or neurological disorder, (b) substance or alcohol abuse/dependence in the past month, (c) heart attack in the past 3 months, (d) pregnancy, (e) presence of contraindications for MRI, (f) presence of contraindications for TMS, such as history of epileptic seizures or certain metal implants^46^, or psychotropic medications that increase risk of seizures, and (g) Quick Inventory of Depressive Symptomatology (16-item, QIDS) self-report questionnaire score of 11 or higher indicating moderate depression^47,48^. All participants completed an MRI pre-examination screening form provided by the Richard M. Lucas Center for Imaging at Stanford University to ensure participant safety prior to entering the MRI scanner. Eligible participants were scheduled for two study visits: an anatomical MRI scan on the first visit and a TMS-EEG session on the second visit. MRI was performed on a GE DISCOVERY MR750 3T MR system (General Electric, Boston, Massachusetts) using a 32 channel head coil. T1 structural scans were acquired using a BRAVO pulse sequence (T1-weighted, sagittal slice thickness 1 mm, acquisition matrix 256 × 256, TR 8 ms, TE 3 ms, FA 15°).

### 2.2. Transcranial Magnetic Stimulation

#### TMS targeting and calibration

TMS was delivered using a MagVenture Cool-B65 A/P figure-of-eight coil from a MagPro X100 system (MagVenture, Denmark). TMS pulse triggering was automated to ensure correct timing in relation to the musical beats, using the MAGIC toolbox for MATLAB^49,50^. Neuronavigation (Localite TMS Navigator, Alpharetta, GA) using each participant’s MRI and a TMS-Cobot-2 system (Axilum Robotics, France) were used to automatically maintain TMS coil placement relative to the participant’s head.

#### Resting motor threshold

To obtain resting motor threshold (rMT), single pulses of TMS were delivered to the hand region of the left M1 with the coil held tangentially to the scalp and at 45° from the midsagittal plane^51–53^. The optimal motor hotspot was defined as the coil position from which TMS produced the largest and most consistent MEP in a relaxed first dorsal interosseous (FDI) muscle^53^. This hotspot was used for both rMT and for iTBS conditions. rMT was determined to be the minimum intensity that elicited an MEP of at least 50 µV peak-to-peak amplitude in relaxed FDI in ≥ 5/10 stimulations^54,55^.

#### MEP assessment

At baseline and at each post-iTBS timepoint (0, 15, 30, and 45 minutes), MEPs were evoked using standard single pulse TMS applied for 100 trials at 120% of rMT in silence with 3 second interstimulus intervals.

#### se-iTBS

To study whether sensory entrainment increases motor plasticity, two iTBS protocols were used: standard iTBS and the seTMS equivalent, se-iTBS. MEPs were recorded before and at 0, 15, and 30 minutes after each protocol. After the first 10 participants were collected, to capture the timing of return to baseline, a 45 minute time point was added for the last 10 participants. Inter-trial coherence (ITC) trough times were used to design the repetitive protocol, based on our prior work^39^. The se-iTBS protocol has 8 seconds of music, followed by 2 seconds of TMS pulses (ten 50 Hz bursts of three pulses) with the last pulse in each burst at 0 ms, −800 ms, −600 ms, −400 ms, −200 ms (first beat), and 0 ms, −800 ms, −600 ms, −400 ms, −200 ms (second beat) in relation to musical strong beat events (Fig. 1A). This music plus TMS pulse train is applied 20 times in sequence, for a total duration of 3 minutes and 20 seconds. iTBS and se-iTBS were applied at 90% of rMT^56–58^ (% MSO M = 56.4, SD = 6.2, range = 49-68). While the original Huang *et al*. (2005) protocol specifies 80% AMT, we used 90% rMT as it is more practical in clinical populations and produces reliable neuroplastic effects in published iTBS studies^56–58^. Intensity, frequency of triplets, and total number of pulses of TMS were all identical between se-iTBS and iTBS. The standard iTBS protocol was delivered in silence, while se-iTBS was delivered with concurrent music presentation as described above. The order of iTBS conditions was randomized across all participants. A break was applied between the two protocols to allow for MEP size to return to baseline; the average break duration was 106.4 minutes (SD = 7.9, range = 89.0-118.0 minutes). Our primary outcome measure was change in the peak-to-peak amplitude of the MEP at 0, 15, 30, and 45 minutes. MEP amplitude was calculated for each trial and then averaged over the trials for each time point and each iTBS condition. We hypothesized that both the se-iTBS and standard iTBS protocols would increase MEP size from baseline (consistent with plasticity induction) but that the effect of se-iTBS would be greater than standard iTBS (Fig. 1B).

**Figure 1.**
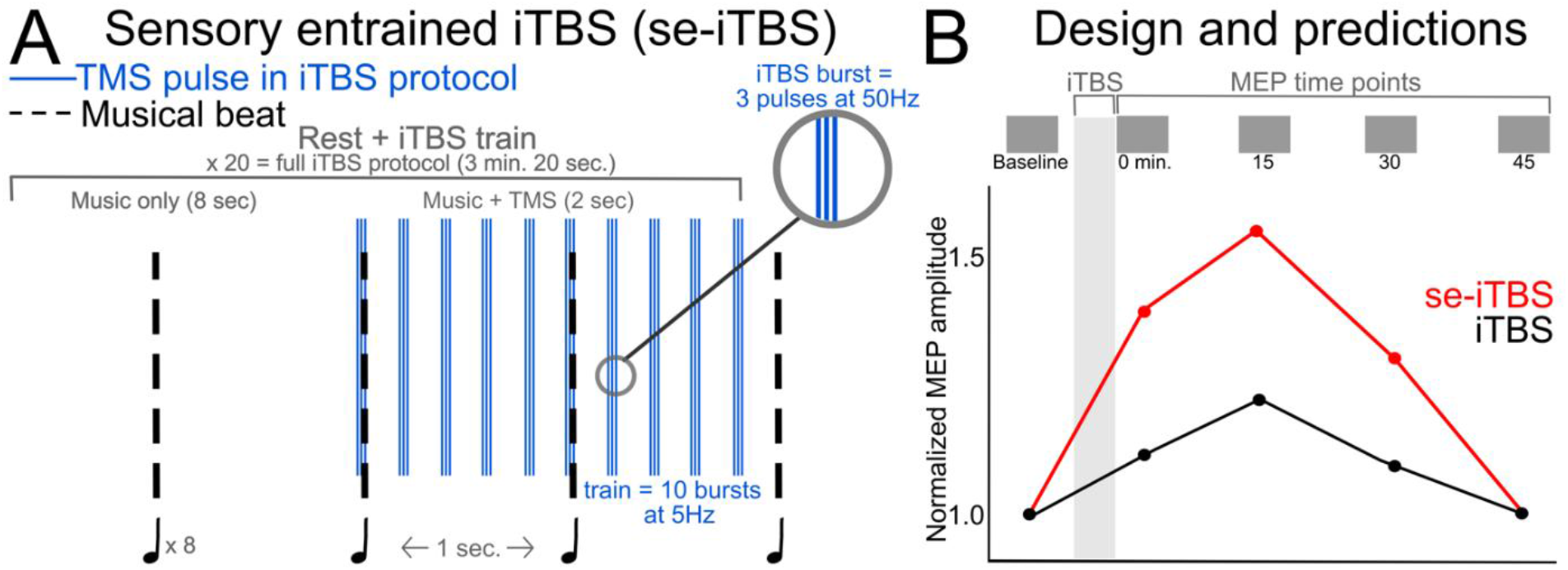
Protocol and Study Design. A) The se-iTBS protocol is identical in timing to the FDA-cleared iTBS protocol except that the participant is listening to music and the protocol bursts align with known maximal excitability states in the brain (ITC troughs in mu, circled in gray), induced using musical beats. The gray trace is an illustration of the expected ITC timeseries. Blue lines are TMS pulse times. Musical sounds were played through earbud-earplugs and noise minimizing over-the-ear muffs were worn to reduce perception of TMS sounds. B) Motor-evoked potentials (MEPs) were recorded from the FDI muscle using single pulse TMS before and after each protocol (se-iTBS, standard iTBS). The order of protocols was randomized across participants.

### 2.3. Auditory stimuli

Auditory stimuli were presented using earbuds at the maximum volume comfortable for each participant. These earbuds are also designed to be earplugs with a Noise Reduction Rating of 25 dB (Elgin USA Ruckus Earplug Earbuds, Arlington, Texas), intended to dampen the TMS “click” sound before reaching the ear canal. For additional dampening of the TMS “click” sound, we used over-the-ear noise-reducing foam-filled earmuffs (3M Ear Peltor Optime 105 behind-the-head earmuffs, NRR 29 dB, Maplewood, Minnesota).

Musical samples used for seTMS were duple or quadruple meter (even groupings of musical beats) and had a tempo of 120 beats per minute (BPM)^44^. This tempo was selected as it falls within the optimal range for beat perception and sensorimotor entrainment (0.5-2 Hz)^59–61^. We selected three musical samples from the Groove Library with high groove ratings (Table S2) to ensure maximal predictive sensory and neural engagement with the musical beats^41,62–66^. Due to alternating strong and weak beat patterns, this tempo results in strong beats ~once per second (1 Hz). We also used a 120 BPM auditory metronome with alternating strong and weak beat sounds (weak = 1/10 amplitude) that has been shown to induce the same excitability dynamics^39^. The auditory metronome consists of 262 Hz tones (middle C), with each tone lasting 60 ms and having a 10 ms duration rise and fall, generated using MATLAB. Like the music, the metronome has strong beats once per second. All auditory stimuli were 30 seconds in length. Seven sounds were used for each participant, selected randomly from all possible sounds and played consecutively for a total duration of 3.5 minutes to cover the full duration of the 3 minute and 20 second iTBS protocol. Standard iTBS was delivered in silence.

For the EEG recording during listening prior to each iTBS protocol, we used the auditory metronome only but for 100 strong beats (100 beats at 1Hz = 1.7 minutes). The metronome was used for EEG recordings to provide a standardized stimulus across beats and to eliminate acoustic complexity that could complicate analysis, while preserving the essential rhythmic structure (1 Hz strong beats, predictable temporal structure) that drives mu desynchronization.

### 2.4. Electromyography

Corticospinal excitability was measured using the peak-to-peak amplitude of motor evoked potentials (MEPs) recorded using electromyography (EMG) from the FDI muscle of the right hand. One surface electrode was placed on the belly of the participant’s muscle. A reference electrode was placed on the lateral face of the proximal interphalangeal joint of the same finger as to not restrict movement. A ground electrode was placed on the styloid process of the wrist of the same hand. To obtain optimal EMG signal, the skin under the electrodes was abraded and cleaned and the electrodes were secured with medical tape. MEPs were elicited by applying single-pulse TMS to the region of the left motor cortex that induced MEPs in FDI. Participants were instructed to keep their hands still and remain relaxed with their right hand on their lap for the duration of the experiment.

#### Preprocessing of EMG

All collected EMG data were processed offline using customized automated scripts running in MATLAB. EMG data were baseline corrected by subtracting the mean value from 20 to 5 ms pre-TMS stimulation from the entire elicited signal. This step centers the signal around zero prior to subsequent peak detection and amplitude calculation; since peak-to-peak amplitude is computed as the difference between minimum and maximum voltage values within a fixed post-stimulus window, any constant DC offset is mathematically cancelled and does not affect the amplitude measurement. Next, trials with artifacts such as pre-activation or concurrent muscle activity were identified. To do this, the root mean square (RMS) of the EMG signal from −200 ms pre-TMS pulse to 13 ms post-TMS pulse, omitting −5 to +5 ms to avoid pulse artifact, was calculated. Trials with RMS values greater than 2.5 standard deviations (SD) from the average RMS of the entire block of trials were removed. Trials were excluded in which MEP amplitudes within the 15–40 ms post-stimulus window were less than or equal to 40 µV^67,68^. This approach is consistent with established TMS methodology, wherein a minimum MEP amplitude threshold is used to define valid corticospinal responses^53^, and with recommendations that averaging MEP amplitude across trials including absent responses can systematically bias estimates of corticospinal excitability^69^. Trials in which MEP amplitudes were larger than 5 standard deviations from the mean were excluded as outliers. The average number of MEP trials remaining after cleaning was 77.4 trials (SD = 17.6) for seTMS and 77.4 trials (SD = 14.8) for standard TMS. By condition the average number of MEP trials for seTMS was baseline = 69.6 (SD = 14.3), 0 min. = 86.2 (SD = 14.5), 15 min. = 70.6 (SD = 21.3), 30 min. = 80.2 (SD = 17.0), 45 min. = 83.2 (SD = 13.3) and for standard TMS was baseline = 72.8 (SD = 13.9), 0 min. = 77.4 (SD = 11.2), 15 min. = 83.4 (SD = 12.5), 30 min. = 76.7 (SD = 17.4), 45 min. = 75.8 (SD = 20.0).

#### Analysis of EMG

First, to assess whether each iTBS protocol modulated cortical excitability, MEP amplitudes at each post-iTBS time point were compared to baseline amplitudes using paired samples *t*-tests, conducted separately for se-iTBS and standard iTBS. Significance was corrected for multiple comparisons using a Bonferroni adjustment (0.05 / 8 = 0.006). Then, to test whether se-iTBS produced greater MEP modulation than standard iTBS, we compared MEP change in amplitude from baseline between the two versions of iTBS at each post-iTBS time point (0-, 15-, 30-, 45-min). MEP peak-to-peak amplitudes were normalized to average baseline amplitudes and compared between the two protocols using paired samples *t*-tests in MATLAB. Normalizing to average baseline amplitudes allows for direct comparison of plasticity magnitude between protocols while controlling for inter-individual variability in absolute MEP amplitudes, which is a common approach in iTBS plasticity studies^25^. Paired *t*-tests were selected because this direct comparison at each timepoint was central to our research question and provides clear, interpretable effect sizes (Cohen’s *d*) for each timepoint. These *t*-tests were performed for each of the 4 time points and the significance level of *α* = 0.05 was corrected for 4 comparisons using a Bonferroni adjustment (0.05 / 4 = 0.0125)^70^. Distributional assumptions were examined; as shown in Supplementary Figure S1, the data distributions, 95% confidence intervals, and normal probability plots support the appropriateness of parametric testing. To note, only 10 participants were tested for MEPs at 45-minutes. Due to the small sample sizes across all timepoints (0-30 min. *n*=20, 45 min. *n*=10), nonparametric Wilcoxon signed rank tests were also used at all 4 time points. These tests do not assume normal distribution and provide robustness checks for the parametric tests.

Although the relationship between timepoint and modulation of MEP amplitude is not expected to be linear^71^, a linear mixed-effect (LME) model can still be useful in understanding whether there is an interaction between a main effect of protocol and time point while accommodating the reduced sample size at the 45-minute timepoint^72,73^. The LME model was implemented specifically to: (1) test for protocol × timepoint interactions while accounting for the unbalanced design, and (2) model within-subject repeated measures with appropriate random effects structure. This statistical analysis was performed in R version 4.4.2 (R Core Team, 2013) to test for a main effect of protocol and an interaction effect with timepoint (0-, 15-, 30-, 45-min), followed by post hoc comparisons between timepoints using Bonferroni’s adjustment for pairwise comparisons. In our model, we specified the protocol as a fixed effect to assess their direct impact, while including participants as a random intercept term to account for individual differences. Both protocol and timepoint were treated as categorical. This LME approach allowed for modeling of within-subject effects with incomplete data (*i*.*e*., that the 45-minute time point was only collected in a subset of participants). The 45-minute timepoint was included in the LME analysis because the missing data were absent by design as the first 10 participants were enrolled before this timepoint was added, and the reason for missingness is therefore fully known and unrelated to the outcome, satisfying the conditions under which LME models with unbalanced designs provide valid estimates^72,74^. We computed estimated marginal means (EMMs)^75^ to summarize model values of each fitted LME model at specific factor levels. If data included in an LME are unbalanced, as in our case, the EMMs represent an estimate of the marginal means that we would have observed if the data had been balanced across conditions.

As an exploratory analysis of individual participant modulation of MEP amplitude, we calculated 4 additional characteristics of the sample:

##### (1) Percentage of participants in whom MEP amplitude was modulated with each protocol

To explore how prevalent MEP modulation was in our sample of participants, we calculated the percentage of the sample that demonstrated an increase in MEP size after each of the protocols. When MEP size normalized to baseline is greater than 1, this indicates that MEPs were qualitatively larger at that time point than at baseline. A threshold of >1 was chosen as it captures any increase above each participant’s own pre-protocol baseline. This method already accounts for the cumulative excitability increases produced by repeated single pulse MEP measurements^76^, which would be expected to affect all recording blocks equally. Furthermore, this threshold was applied consistently to both conventional iTBS and se-iTBS, meaning any stimulation-related effects would be incorporated equally in both conditions and would not bias comparisons between them. We acknowledge this threshold may include participants whose MEPs fluctuated near baseline, and that future sham-controlled designs would allow for a more empirically grounded responder criterion. We calculated the percentage of participants who had a normalized MEP of greater than 1 for each of the four post-iTBS timepoints.

##### (2) Percentage of participants with larger MEP modulation after se-iTBS than after iTBS

To explore how often an increase in MEP size from baseline was larger with se-iTBS vs. with standard iTBS, we calculated the percentage of the sample that showed a qualitatively larger normalized MEP after se-iTBS for each of the four post-iTBS timepoints.

##### (3) Time of peak MEP modulation for each protocol

To explore the timing of peak MEP modulation after both protocols we identified for each participant the time point out of the four with the largest normalized MEP and then calculated for each time point the percentage of the sample that peaked at that time point.

##### (4) Time of maximum difference in MEP modulation between the protocols

To explore the time course of MEP modulation between the protocols, we calculated the percentage of participants who had the qualitatively largest absolute difference between protocols in the normalized MEP at each of the four post-iTBS timepoints.

### 2.5. Electroencephalography

To confirm the expected beat-related mu dynamics previously reported^39,44^, EEG was recorded during beat listening without TMS prior to each of the iTBS protocols. During these two recordings (one before each iTBS protocol), participants listened to the auditory metronome for 100 beats with eyes open and without moving their bodies. 64-channel EEG was obtained using a BrainVision actiCHamp Plus amplifier, with ActiCAP slim active electrodes in an extended 10–20 system montage (actiCHamp, Brain Products GmbH, Munich, Germany) with a 25 kHz sampling rate^77^. EEG data were online referenced to Cz and recorded using BrainVision Recorder software v1.24.0001 (Brain Products GmbH, Germany). Impedances were monitored and percentage of channels with impedances <10 kΩ was 99.9 ± SD 0.3% prior to se-iTBS and 99.5 ± SD 0.9% prior to standard iTBS. Electrode locations were digitized using Localite.

#### Preprocessing of EEG

EEG data were pre-processed offline using the Resting-state Semi-Automated Preprocessing pipeline (R-SAP, described below, available at https://github.com/jross4-stanford/R-SAP)^78^ and EEGLab v2021.1 in MATLAB R2021a (Mathworks, Natick, MA, USA).

#### Resting-state Semi-Automated Preprocessing (R-SAP)

Data were epoched and time-locked to each strong beat event, with epochs spanning from −500 ms to +500 ms relative to beat onset. This resulted in 100 epochs per recording. Data were downsampled to 1000 Hz. Low-pass (49 Hz) and high-pass (1Hz) filters were applied using a zero-phase 4th order Butterworth filter. Conservative channel rejection and epoch rejection, and noise removal were applied using the *clean_rawdata* function (FlatlineCriterion = 5, ChannelCriterion = 0.8, BurstCriterion = 5, WindowCriterion = 0.5). Missing/removed channels were interpolated using spherical interpolation, and data were re-referenced to the average. The mean number of channels removed was 0.6 channels (SD = 1.3, range = 0-5) prior to se-iTBS and 0.4 channels (SD = 1.1, range = 0-4) prior to standard iTBS. The mean number of epochs remaining was 98.4 epochs (SD = 5.3, range = 76-100) prior to se-iTBS and 96.4 epochs (SD = 10.5, range = 54-100) prior to standard iTBS. Because recordings were made with 64 channels, and the signals were unlikely to have that many independent sources, PCA was used to reduce dimensionality prior to ICA to 30 dimensions. This approach can improve decomposition^79,80^ and signal to noise ratio of large sources^81^. Fast independent component analysis (FastICA) was run^82^ and the Multiple Artifact Rejection Algorithm (MARA)^83,84^ was used to identify components with high likelihood of being non-brain artifacts (posterior_artifactprob > 0.30). These components were removed, and remaining components were reviewed using the open source TMS-EEG Signal Analyzer (TESA v1.1.0-beta) extension for EEGLAB^85,86^ (http://nigelrogasch.github.io/TESA/), allowing for additional components to be rejected by an expert reviewer if necessary. Mean number of components remaining after cleaning was 10.6 components (SD = 2.5, range = 7-15) prior to se-iTBS and 11.8 components (SD = 3.7, range = 6-18) prior to standard iTBS. The proportion of ICs rejected is consistent with the rate reported using the same pipeline in prior published work^44,78^ and is expected for high-density 64-channel EEG recordings, for which the majority of ICs are typically non-brain in origin^87^.

#### Analysis of EEG

To verify that passive listening to the auditory metronome induced mu desynchronization (low ITC) prior to the beat time^44^, time-frequency analysis was completed for each participant at each channel and then averaged across three channels from over the left motor cortex (C5, C3, C1). These electrodes are positioned over the primary motor cortex for the right hand and are standard scalp electrode locations for recording sensorimotor mu rhythms^88–92^. Time-frequency features were computed with the *newtimef* function in EEGLAB^93^ using linear spaced Morlet wavelets between 6 and 48 Hz with a fixed window size of 500 ms resulting in 3 cycles at the lowest frequency of 6 Hz. Log mean baseline power spectrum between −500 and −200 ms preceding beat times was removed^94–96^. The 500 ms window size was chosen to ensure that the time–frequency representation from each individual stimulus was not contaminated by either of the surrounding stimuli, which were 1000 ms apart. These computations were used to determine the event-related spectral ITC^93^. ITC is calculated by extracting the phase angle at each time–frequency point for each trial and comparing the phase angles across trials for coherence. This provides a coherence measure between 0 and 1, where 1 indicates complete coherence across trials for a given time–frequency point, and 0 indicates no coherence across trials. Alpha ITC was extracted by averaging the power at each frequency bin between 8 and 14 Hz^88,97^. The largest ITC trough was expected to occur ~200 ms prior to beat times^44^, so ITC troughs were calculated as the local minima between −222 and −99 ms, for each individual participant.

ITC troughs were compared before se-iTBS and iTBS using paired samples *t*-tests. Because the order of plasticity protocols was randomized across participants, we also asked whether there were differences between the two EEG recordings in the day (first vs. second recording), using paired samples *t*-tests and nonparametric Wilcoxon signed rank tests.

## 3. Results

### 3.1. Modulation of MEP Amplitude

We first assessed whether each iTBS protocol modulated MEP amplitudes relative to baselines. Overall, MEPs were significantly larger after the iTBS protocols at at least one time point than at baseline, supporting that both protocols modulated the MEP amplitude and that these changes were in the same direction (increase from baseline). Specifically, after correcting for the multiple comparisons^70^, MEPs were larger after se-iTBS at the first three time points (0-min *t*(19) = −6.82, *p* = 0.000002, Cohen’s *d* = −1.52, 15-min *t*(19) = −4.29, *p* = 0.0004, *d* = −0.96, and 30-min *t*(19) = −4.23, *p* = 0.0004, *d* = −0.95) but not at 45-minutes (*t*(9) = −1.54, *p* = 0.16), and after standard iTBS at the first time point (0-min *t*(19) = −4.18, *p* = 0.0005, *d* = −0.94) but not the last three time points (15-min *t*(19) = −2.47, *p* = 0.02, and 30-min *t*(19) = −1.78, *p* = 0.09, 45-min *t*(9) = 0.46, *p* = 0.65). See Table S3 for all test statistics, means, standard deviations, and standard errors of the means. The average percent increase from baseline in MEP size at 0-min for se-iTBS was 54.58% (15-min 42.17%, 30-min 36.67%, 45-min 16.79%) and for iTBS was 25.97% (15-min 19.34%, 30-min 16.66%, 45-min 4.87%, Table S4). For all individual participant MEPs, see Supplementary Figures S5-S24. Baseline MEPs were not significantly different between se-iTBS and iTBS (Figure S2, *t*(19) = −0.26, *p* = 0.80, Cohen’s *d* = 0.06, See tables S7-8 for parametric and nonparametric comparisons) and were not predicted by age of the participants (*R*^*2*^ = 0.00068, *F*(1,17) = 0.011, *p* = 0.92).

### 3.2. Comparison of MEP Modulation Between Protocols

We next tested whether se-iTBS produced greater MEP modulation than standard iTBS. By normalizing MEP size to baselines, we compared MEP change in amplitude (hereafter referred to as MEP modulation) between the two versions of iTBS at each post-iTBS time point (0-, 15-, 30-, 45-min), correcting for multiple comparisons^70^. se-iTBS induced more MEP modulation than standard iTBS across time points (Figure 2). Specifically, se-iTBS induced significantly more MEP modulation than standard iTBS at the first three time points (0-min *t*(19) = 3.98, *p* = 0.0008, Cohen’s *d* = 0.89, 15-min *t*(19) = 4.29, *p* = 0.0004, *d* = 0.96, and 30-min *t*(19) = 2.91, *p* = 0.009, *d* = 0.65), and there was no difference between protocols at 45-minutes (*t*(9) = 2.20, *p* = 0.06). MEP modulations were not predicted by age of participants (0 min: *R*^*2*^ = 0.0068, *F*(1,17) = 0.12, *p* = 0.74; 15 min: *R*^*2*^ = 0.00088, *F*(1,17) = 0.015, *p* = 0.90; 30 min: *R*^*2*^ = 0.041, *F*(1,17) = 0.73, *p* = 0.40). se-iTBS increased MEP modulation by an average of 17.59% compared to standard iTBS (0-min 22.71%, 15-min 19.13%, 30-min 17.15%, 45-min 11.37%, Table S5). See Table 1 for all test statistics and protocol means, standard deviations, and standard errors of the means. Due to the relatively small sample sizes across all time points (*n*=20, *n*=10), nonparametric Wilcoxon signed rank tests were also used and showed that MEP modulation from se-iTBS was greater than from standard iTBS at all 4 time points (see Table S6). See Supplementary Figure S1 for data distributions, 95% confidence intervals, and normal probability plots. In summary, direct comparisons between the two versions of iTBS, using parametric and nonparametric methods, showed that se-iTBS induced more modulation of MEP amplitude than standard iTBS.

**Table 1.**
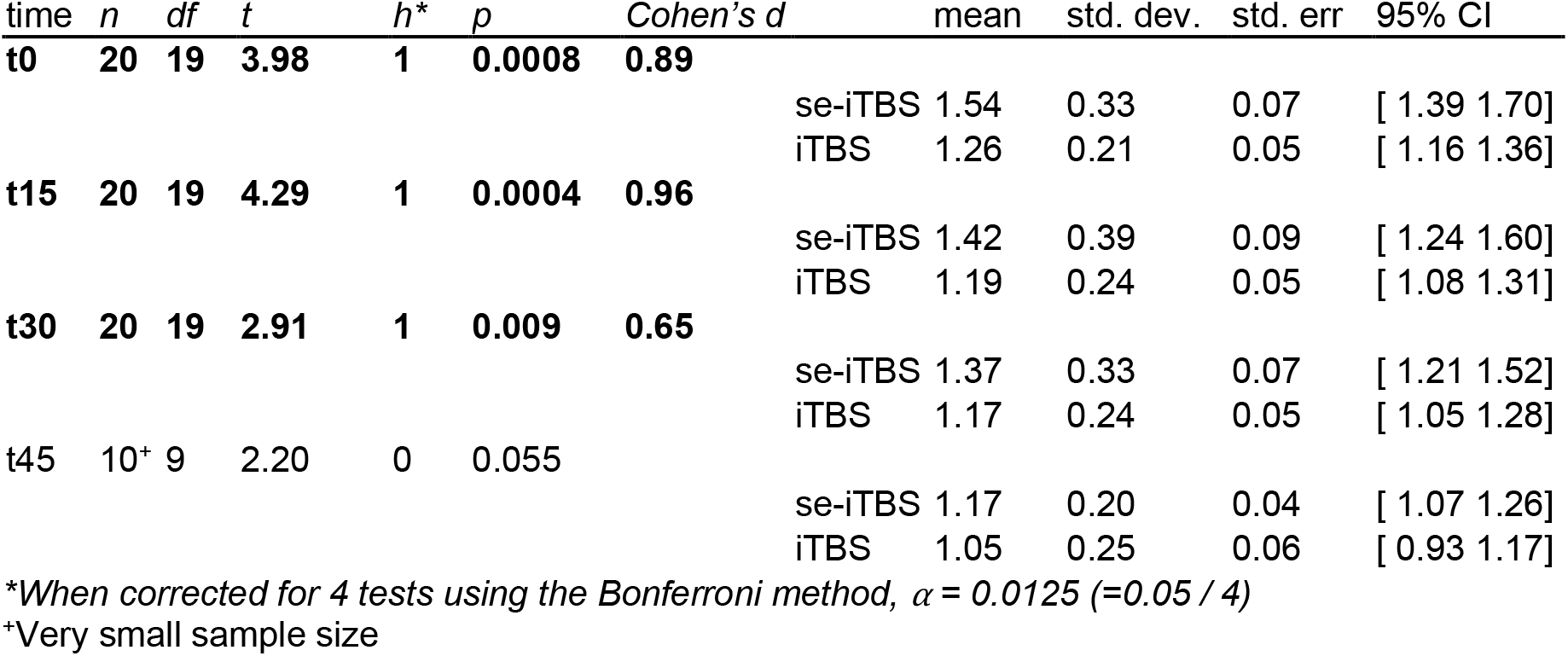
Comparison of MEP modulation between protocols. Paired t-tests at each time point.

**Figure 2.**
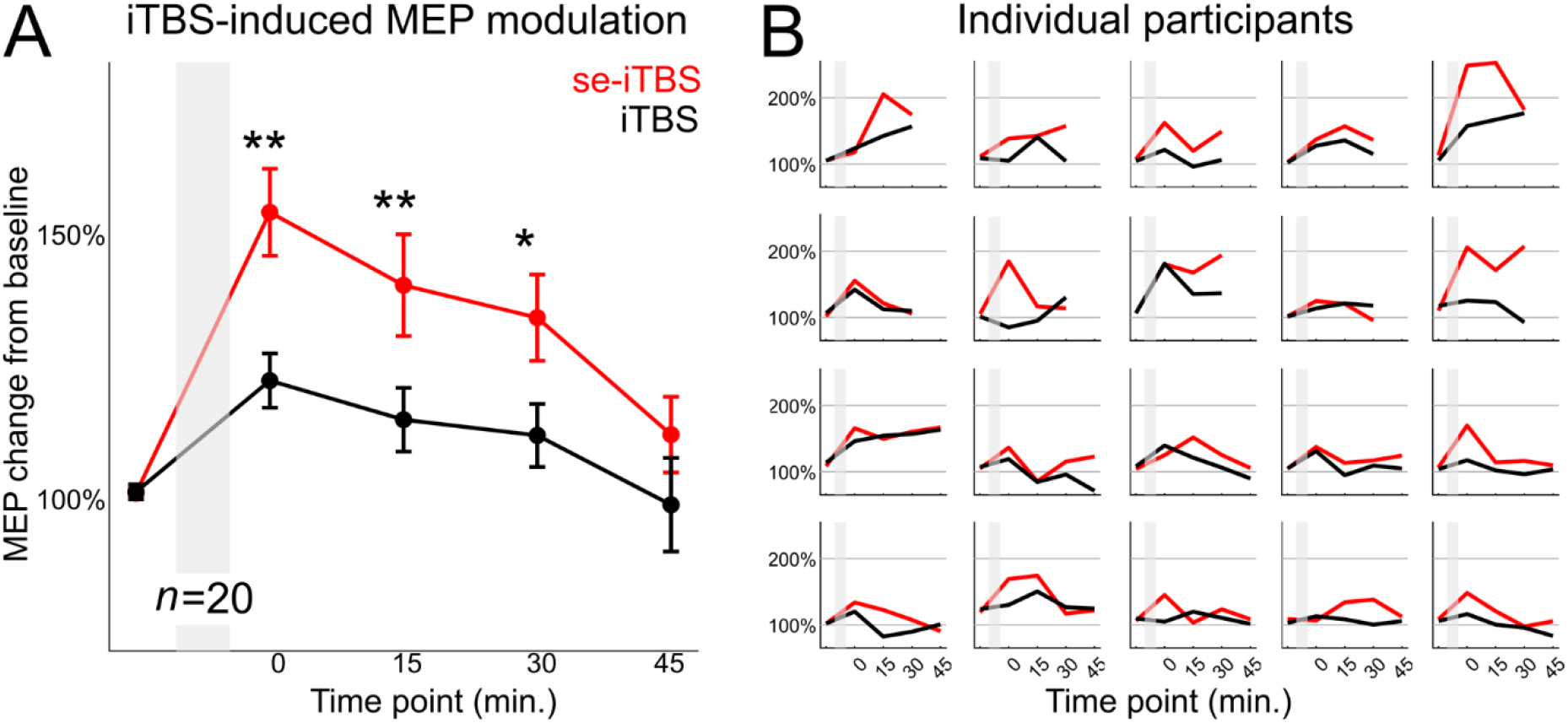
se-iTBS induces more MEP modulation than standard iTBS. (A) MEPs were averaged over all participants (*n*=20), with standard error of the mean shown across participants. The shaded region is when iTBS was applied. se-iTBS induced more MEP modulation than standard iTBS at the first three time points (0-min *t*(19) = 3.98, *p* = 0.0008, Cohen’s *d* = 0.89, 15-min *t*(19) = 4.29, *p* = 0.0004, *d* = 0.96, and 30-min *t*(19) = 2.91, *p* = 0.009, *d* = 0.65). ** *p* < 0.001, * *p* < 0.01. An LME was additionally used to test for a protocol by time point interaction, which was not found (F(3,113) = 0.87, *p* = 0.46), but main effects were significant of protocol (F(1,113) = 44.95, *p* < 0.0001) and time point (F(3,113) = 5.70, *p* = 0.001). (B) Individual participants’ MEP modulations.

Next, we used a linear mixed-effect model to test for a protocol by time point (0-, 15-, 30-, 45-min) interaction (see Methods for more details). The model revealed significant effects of protocol (F(1,113) = 44.95, *p*<.0001) and of time point (F(3,113) = 5.70, *p*=.001) but no protocol by time point interaction (F(3,113) = 0.87, *p*=.46). Post-hoc pairwise comparisons with Bonferroni correction indicated significant differences between time points 0 min and 30 min (*t*(113) = 3.12, *p* = 0.01), and between time points 0 min and 45 min (*t*(113) = 3.72, *p* = 0.002). These post-hoc results suggest that the MEP effects reduced with time after the protocol. Because we did not observe a protocol by time point interaction, we can interpret that this reduction in MEP effects with time did not depend on which protocol was used. See Table 2 for test statistics for all post-hoc comparisons, and Figure S3 for LME and EMMs. We found no systematic changes in MEP modulation with iTBS intensity in percentage of maximum stimulator output (Supplementary Figure S4) at 0-min (se-iTBS *R*^*2*^ = 0.11, *F*(2,18) = 1.15, *p* = 0.34; iTBS *R*^*2*^ = 0.0012, *F*(2,18) = 0.011, *p* = 0.99), 15-min (se-iTBS *R*^*2*^ = 0.074, *F*(2,18) = 0.72, *p* = 0.50; iTBS *R*^*2*^ = 0.050, *F*(2,18) = 0.48, *p* = 0.63), 30-min (se-iTBS *R*^*2*^ = 0.18, *F*(2,18) = 2.06, *p* = 0.16; iTBS *R*^*2*^ = 0.0031, *F*(2,18) = 0.028, *p* = 0.97), or 45-min (se-iTBS *R*^*2*^ = 0.012, *F*(2,8) = 0.048, *p* = 0.95; iTBS *R*^*2*^ = 0.0024, *F*(2,8) = 0.0098, *p* = 0.99).

**Table 2.** Comparison of MEP modulation between protocols and across time points. Linear mixed-effect model and post-hoc comparisons.

| factor | df | F-value | h | p |
| --- | --- | --- | --- | --- |
| **protocol** | **1,113** | **44.95** | **1** | **<0.0001** |
| **time point** | **3,113** | **5.70** | **1** | **0.001** |
| protocol* time point | 3,113 | 0.87 | 0 | 0.46 |

| contrasts | df | t | h | p* |
| --- | --- | --- | --- | --- |
| 0min-15min | 113 | 2.18 | 0 | 0.19 |
| **0min-30min** | **113** | **3.12** | **1** | **0.01** |
| **0min-45min** | **113** | **3.72** | **1** | **0.002** |
| 15min-30min | 113 | 0.94 | 0 | 1.00 |
| 15min-45min | 113 | 2.02 | 0 | 0.28 |
| 30min-45min | 113 | 1.29 | 0 | 1.00 |

\*Corrected for 6 tests using the Bonferroni method

### 3.3. Individual Participants

Individual participant MEP modulations for each protocol are shown in Figure 3A. To learn more about MEP modulations in individual participants, we explored four additional characteristics of the sample (Figure 3B-C):

**Figure 3.**
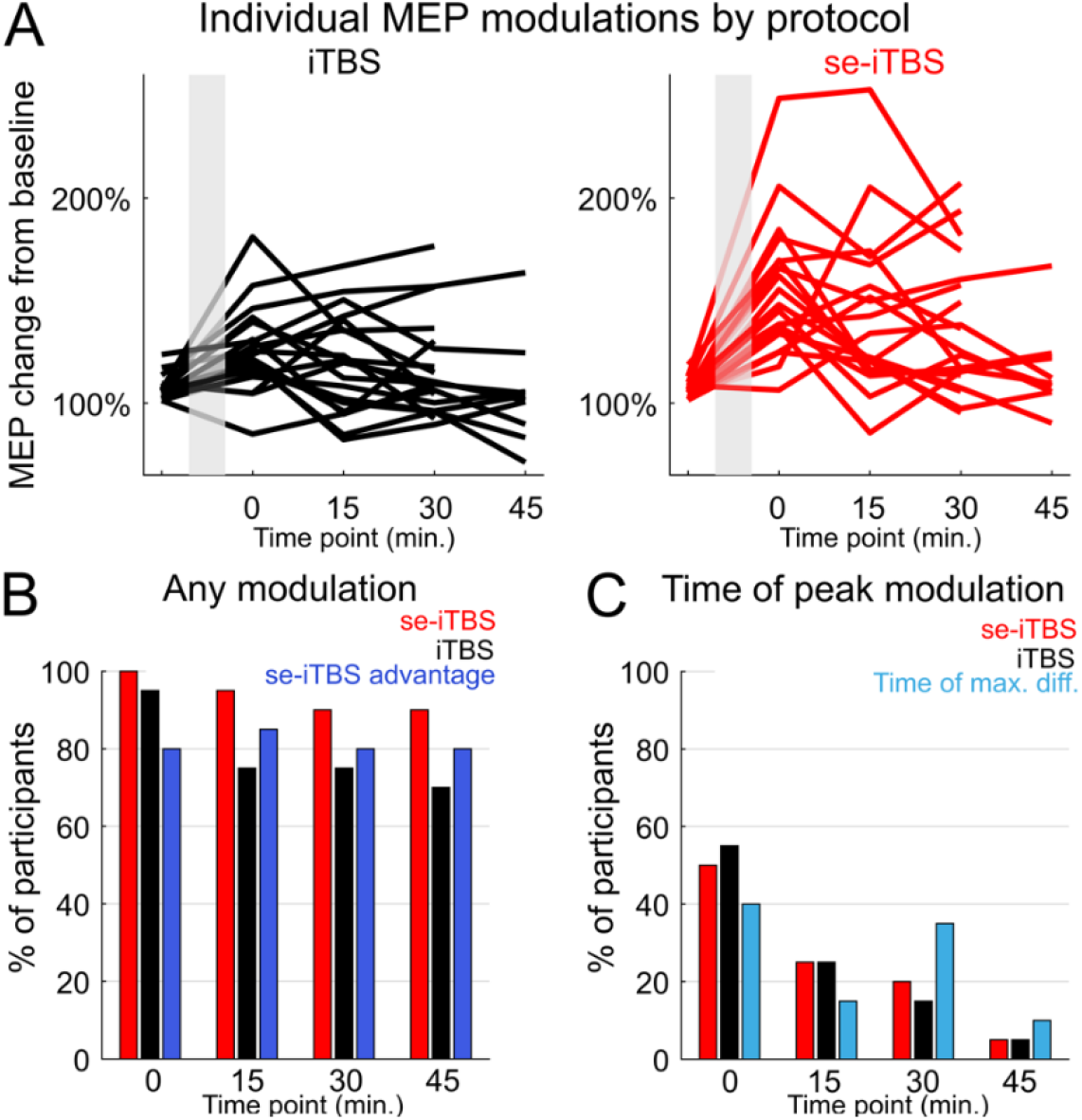
iTBS induced robust individual participant MEP changes. (A) Individual participants’ MEP modulations by protocol. The shaded region is when iTBS was applied. (B) Percentage of participants with their peak MEP change at each time point (red, black) and with their maximum difference in MEP modulation between iTBS protocols at each time point (light blue). (C) Percentage of participants in which any MEP modulation was induced with each iTBS protocol (red, black) and the percentage of participants who had more MEP modulation with se-iTBS than with iTBS (dark blue).

#### (1) Percentage of participants in whom MEP amplitude was modulated with each protocol

For se-iTBS, the percentage of participants showing larger MEPs than at baseline at 0-min was 100%, at 15-min was 95%, at 30 min was 90%, and at 45-min was 90%. For iTBS, the percentage of participants showing larger MEPs than at baseline at 0-min was 95%, at 15-min was 75%, at 30-min was 75%, and at 45-min was 70% (Figure 3B).

#### (2) Percentage of participants with larger MEP modulation after se-iTBS than after iTBS

at 0-min was 80%, at 15-min was 85%, at 30-min was 80%, and at 45-min was 80% (Figure 3B dark blue).

#### (3) Time of peak MEP modulation for each protocol

For se-iTBS, peak MEP change occurred at 0-min in 50% of participants, at 15-min in 25% of participants, at 30-min in 20% of participants, and at 45-min in 5% of participants. For iTBS, peak MEP change occurred at 0-min in 55% of participants, at 15-min in 25% of participants, at 30-min in 15% of participants, and at 45-min in 5% of participants (Figure 3C).

#### (4) Time of maximum difference in MEP modulation between the protocols

the greatest difference in MEP modulation between the two protocols was at 0-min in 40% of the participants, at 15-min in 15% of the participants, at 30-min in 35% of the participants, and at 45-min in 10% of the participants (Figure 3C light blue).

### 3.4. Electroencephalography

Having demonstrated that se-iTBS produces enhanced MEP modulation compared to standard iTBS, we next sought to validate the underlying mechanism of our approach. Our se-iTBS method relies on the premise that auditory rhythms induce predictable mu desynchronization patterns that can be leveraged to optimize TMS timing. To confirm this foundational assumption in our participants, we assessed EEG mu desynchronizations during passive listening to the auditory metronome used in our protocol. ITC troughs in mu occurred during passive listening to the metronome before both se-iTBS and iTBS. Paired samples *t*-tests confirmed that trough latency did not differ significantly between conditions (Table S9, *t*(19) = −1.39, *p* = 0.18; Cohen’s *d* = 0.31, pre-se-iTBS M = −172.05 ms, SD = 44.44; pre-iTBS M = −151.70 ms, SD = 42.40). Because the order of iTBS protocols was randomized across participants, we also asked whether there were differences between the two EEG recordings in the day (first vs. second recording). Trough latency did not differ significantly between the first and second recording (Table S9, *t*(19) = −0.36, *p* = 0.72; Cohen’s *d* = 0.08, first recording M = −164.60 ms, SD = 43.32; second recording M = −159.15 ms, SD = 45.81). Wilcoxon signed rank tests also support no difference between pre-se-iTBS vs. pre-iTBS and between first recording vs. second recording (see Table S10 for statistics). In summary, our data from passive listening to the auditory metronome showed induced mu desynchronization prior to the beats that is consistent with prior literature^38,39,44^ and did not differ between the two recordings in the day (Figure S2).

## 4. Discussion

### 4.1. Summary of findings

In this study, we present a novel approach called Sensory Entrained iTBS (se-iTBS) that uses music to synchronize excitability to prime the brain for stimulation. We applied se-iTBS to M1 and in a within-subjects crossover design compared effects with standard iTBS using MEPs as a measure of corticospinal excitability. Our key findings were: 1) Both versions of iTBS induced corticospinal modulations of excitability; 2) se-iTBS resulted in larger effects than standard iTBS. se-iTBS increased MEP amplitudes significantly more than standard iTBS for at least 30 minutes. This led to more than double the size of the modulation (55% vs. 26% change in MEP amplitude; Figures 1-2). 3) se-iTBS resulted in more consistent effects across individuals than standard iTBS. The enhanced effect from se-iTBS was observed in 80-85% of individual participants (Figure 3). Together, this work demonstrates that sensory entrained iTBS increased corticospinal modulation.

### 4.2. Need for optimization of iTBS to primary motor cortex

Despite its emerging therapeutic use, fundamental gaps in motor cortex iTBS protocols urgently need addressing, as individual response variability undermines its therapeutic potential. While iTBS has shown promise in inducing corticospinal plasticity and improving upper limb function in stroke patients^12^, its effects are inconsistent, with some studies reporting significant variability in iTBS-induced change in MEP amplitudes^22,98^. iTBS effects on the motor cortex are influenced by factors such as baseline corticospinal excitability, M1 mu oscillations^24^, and precise neuroanatomical targeting^23^. Furthermore, even though preclinical evidence supports the idea that iTBS can induce lasting structural changes in the spinal cord and enhance corticospinal motor output^3^, more research is needed to optimize iTBS parameters across different applications^20^. Together these findings highlight the need for more refined protocols that can reliably enhance excitability and improve clinical outcomes. Our results directly address this challenge by showing that se-iTBS achieved consistent MEP enhancement in over 80% of participants, more than doubling the effects of standard iTBS (Figure 3). This suggests that se-iTBS could mitigate the variability observed in traditional iTBS protocols by leveraging endogenous neural oscillations, underscoring the potential of incorporating brain-state-dependent neuromodulation to optimize iTBS. The consistent enhancement across participants suggests that synchronizing iTBS with endogenous neural oscillations represents a promising approach to address the variability that has limited traditional iTBS applications. By leveraging sensorimotor oscillations, se-iTBS offers a practical framework that bridges fundamental electrophysiological mechanisms with improved clinical applicability, potentially expanding the therapeutic utility of iTBS across neurological and psychiatric conditions. Advances in research on optimal iTBS parameters for inducing lasting structural and functional changes in the motor system should be incorporated into future se-iTBS research.

### 4.3. Neural mechanisms underlying seTMS

Musical rhythms provide a powerful yet unexplored gateway to control cortical excitability and enhance plasticity. This is because musical beat perception involves complex excitability dynamics^31–33,38–40,59,61,99–101^. These excitability dynamics have often been described as sensorimotor coupling, or “covert action”, and occur without overt body movement^31–33,38– 40,59,61,99,101–103^. These dynamics appear to be essential for accurate rhythm perception^61,99,103,104^, as demonstrated by both neuroimaging studies and cases of impaired perception following brain lesions^105–112^. Neurophysiological studies using MEG and EEG have revealed beat-related excitability dynamics across multiple brain regions^31–33,37–40^, including auditory, sensorimotor, premotor, parietal, occipital and frontal cortices, as well as the cerebellum. Importantly, these dynamics reflect the beat times that were predicted by the participant, suggesting top-down modulation of auditory perception^33^ and appear to be relevant to TMS-related excitability^41,44^. The 1 Hz tempo of our musical stimuli was specifically chosen to fall within the optimal range for beat perception and sensorimotor entrainment, inducing the mu desynchronization patterns that underlie our approach^31–33,38–40,44^. These findings align with clinical evidence showing that auditory-motor entrainment can effectively modulate neural circuits to improve motor function^42,43^. Our findings significantly extend this understanding by demonstrating that TMS pulses delivered at optimal mu desynchronization timing just before musical beats significantly enhances MEP modulations (Figure 2). Further, the consistency of these effects across individual participants (Figure 3) suggests that musical rhythms provide a reliable method for targeting windows of optimal cortical excitability. Our results contribute to this growing literature in establishing a mechanistic framework for enhancing neuromodulation through brain-state alignment.

### 4.4. MEP modulation and plasticity considerations

Our use of the term “plasticity” in this manuscript refers to the modulation of MEP amplitudes following iTBS, rather than direct evidence of specific cellular or synaptic mechanisms. While iTBS is commonly described as a plasticity protocol in the literature, the precise neural mechanisms underlying MEP modulation remain incompletely understood and may involve various forms of neural adaptation including synaptic plasticity, changes in cortical excitability, changes in brain-states, or altered network dynamics. The sustained nature of the MEP changes we observed (lasting at least 30 minutes) suggests engagement of mechanisms beyond transient excitability shifts, consistent with plasticity-like processes. However, definitive characterization of the underlying mechanisms would require additional investigations using techniques such as paired-pulse TMS protocols, pharmacological interventions, or invasive direct neural recordings^113,114^. Our use of “plasticity” should therefore be interpreted as a functional description of the sustained neural modulation rather than a claim about specific molecular or cellular mechanisms.

## 5. Limitations and future directions

While our study demonstrates the potential of se-iTBS to enhance plasticity, several limitations should be addressed in future research. Our investigation was limited to healthy participants in a controlled laboratory setting; future studies should evaluate se-iTBS effects in clinical populations where enhanced plasticity could be therapeutically beneficial, such as stroke recovery or depression. Our study used a standardized set of auditory stimuli; future work should investigate whether personalizing the sound selection could further enhance iTBS modulation effects. In addition to personalized music, future work should systematically investigate how specific acoustic features^115–117^ optimize entrainment for enhanced neuromodulation. Future work should also test iTBS paired with a pure 1 Hz tone to isolate the specific contribution of musical complexity beyond simple isochronous beats. A direct comparison of ITC trough dynamics between musical samples and metronomes should be conducted in future studies, to confirm whether either stimulus can be used interchangeably for quantifying ITC troughs in the context of se-iTBS delivery. We only tested a fixed se-iTBS timing across participants; systematically varying the timing in relation to auditory beats could reveal optimal parameters for enhancing MEP changes. Specifically, testing iTBS delivery at different temporal relationships to the auditory beats would help determine whether the enhanced effects depend on precise temporal alignment or simply the presence of auditory rhythms, further elucidating the role of temporal synchronization in optimizing neuroplastic responses. A related approach would be to record EEG prior to iTBS delivery to obtain individually-measured ITC troughs that could provide more precise temporal alignment of stimulation to each participant’s neural oscillations and represents a practical avenue for future optimization. We only examined effects up to 45 minutes post-stimulation in a subset of participants; longer follow-up periods are needed to fully characterize the duration of enhanced MEP modulation. A further limitation is that the two protocols were administered in the same session. Although protocol order was counterbalanced across participants and MEP amplitudes returned to baseline prior to the second protocol, we cannot rule out that the first protocol may have subtly influenced the neural substrate available for plasticity induction during the second. Future studies should administer se-iTBS and standard iTBS on separate days to fully isolate each protocol’s effects. Moving forward, key directions for research include examining 1) whether se-iTBS can increase behavioral effects and clinical scales, 2) whether brain regions outside of M1 show enhanced modulations with se-iTBS, particularly those relevant to psychiatric treatments^21,113,114^, 3) the neural mechanisms underlying the enhanced iTBS effects through combined TMS-EEG recordings during stimulation, 4) if individual factors such as musical training or musical preferences can predict iTBS effects or be used to optimize the effects. We will also 5) develop optimized se-iTBS protocols that maximize modulations while maintaining safety and tolerability, 6) systematically investigate how different types of auditory stimuli and timing relationships between sounds and TMS pulses affect iTBS modulations, and 7) employ single-protocol designs on separate days with extended post-stimulation follow-up beyond 45 minutes to more definitively characterize the independent and sustained effects of each protocol. These directions could significantly advance our understanding of rhythm-enhanced neuromodulation and accelerate translation to clinical applications.

## 6. Conclusions

This study introduces Sensory Entrained iTBS (se-iTBS), a novel approach that leverages sensorimotor oscillations using auditory rhythms to enhance iTBS effects. Our findings demonstrate that se-iTBS more than doubles the corticospinal neuroplastic effects of standard iTBS, producing significant enhancement in over 80% of participants. se-iTBS offers a readily implementable, low-cost method for optimizing TMS outcomes with potential applications in basic neuroscience and clinical treatment across neurological and psychiatric disorders.

## Acknowledgments

We would like to acknowledge the contributions of all our research participants. We extend gratitude to the members of the Precision Neurotherapeutics Laboratory for helpful feedback on the article and throughout the course of the study.

## Author contributions

JMR, JG, UH, CCC, SP, NFC, TF, SM, APL, and CJK conceptualized and designed the study. JMR and CJK acquired funding. JMR and JT programmed the experiment. JMR, JT, LF, and JWH collected the data. JMR, LF, JG, JWH, NFC, and SP conducted the analyses. All authors interpreted the results. All authors contributed to the writing of the manuscript. All authors provided intellectual contributions to and approval of the final manuscript.

## Availability of Data and Materials

The datasets generated and/or analyzed during the current study are available upon request.

## Supplementary Information

The online version contains supplementary material.

## References

1. Di Lazzaro V, Pilato F, Dileone M, et al. The physiological basis of the effects of intermittent theta burst stimulation of the human motor cortex. J Physiol. 2008;586(16):3871–3879.

2. Huang YZ, Edwards MJ, Rounis E, Bhatia KP, Rothwell JC. Theta Burst Stimulation of the Human Motor Cortex. Neuron. 2005;45(2):201–206.

3. Amer A, Martin JH. Repeated motor cortex theta-burst stimulation produces persistent strengthening of corticospinal motor output and durable spinal cord structural changes in the rat. Brain Stimulation. 2022;15(4):1013–1022.

4. Hanlon CA, Smith HR, Epperly PM, Collier M, Galbo LK, Czoty PW. Priming the pump? Evaluating the effect of multiple intermittent theta burst sessions on cortical excitability in a nonhuman primate model. Brain Stimulation. 2022;15(3):676–677.

5. Yesavage JA, Fairchild JK, Mi Z, et al. Effect of Repetitive Transcranial Magnetic Stimulation on Treatment-Resistant Major Depression in US Veterans: A Randomized Clinical Trial. JAMA Psychiatry. 2018;75(9):884.

6. Madore MR, Kozel FA, Williams LM, et al. Prefrontal transcranial magnetic stimulation for depression in US military veterans – A naturalistic cohort study in the veterans health administration. J Affect Disord. 2022;297:671–678.

7. Blumberger DM, Vila-Rodriguez F, Thorpe KE, et al. Effectiveness of theta burst versus high-frequency repetitive transcranial magnetic stimulation in patients with depression (THREE-D): a randomised non-inferiority trial. Lancet. 2018;391(10131):1683–1692.

8. Chail A, Saini RK, Bhat PS, Srivastava K, Chauhan V. Transcranial magnetic stimulation: A review of its evolution and current applications. Ind Psychiatry J. 2018;27(2):172–180.

9. Bai Z, Zhang JJ, Fong KNK. Immediate Effects of Intermittent Theta Burst Stimulation on Primary Motor Cortex in Stroke Patients: A Concurrent TMS-EEG Study. IEEE Trans Neural Syst Rehabil Eng. 2023;31:2758–2766.

10. Ding Q, Chen S, Chen J, et al. Intermittent Theta Burst Stimulation Increases Natural Oscillatory Frequency in Ipsilesional Motor Cortex Post-Stroke: A Transcranial Magnetic Stimulation and Electroencephalography Study. Front Aging Neurosci. 2022;14:818340.

11. Cao Z, Zhang J, Lu Z, et al. Physical Activity, Mental Activity, and Risk of Incident Stroke: A Prospective Cohort Study. Stroke. 2024;55(5):1278–1287.

12. Huang W, Chen J, Zheng Y, et al. The Effectiveness of Intermittent Theta Burst Stimulation for Stroke Patients With Upper Limb Impairments: A Systematic Review and Meta-Analysis. Front Neurol. 2022;13:896651.

13. Yang Q, Xu S, Chen M, et al. Effects of the Left M1 iTBS on Brain Semantic Network Plasticity in Patients with Post-Stroke Aphasia: A Preliminary Study. J Integr Neurosci. 2023;22(1):24.

14. Lefaucheur JP, Aleman A, Baeken C, et al. Evidence-based guidelines on the therapeutic use of repetitive transcranial magnetic stimulation (rTMS): An update (2014–2018). Clin Neurophysiol. 2020;131(2):474–528.

15. O’Reardon JP, Solvason HB, Janicak PG, et al. Efficacy and safety of transcranial magnetic stimulation in the acute treatment of major depression: a multisite randomized controlled trial. Biol Psychiatry. 2007;62(11):1208–1216.

16. Cole EJ, Stimpson KH, Bentzley BS, et al. Stanford Accelerated Intelligent Neuromodulation Therapy for Treatment-Resistant Depression. Am J Psychiatry. 2020;177(8):716–726.

17. Cole EJ, Phillips AL, Bentzley BS, et al. Stanford Neuromodulation Therapy (SNT): A Double-Blind Randomized Controlled Trial. Am J Psychiatry. 2022;179(2):132–141.

18. Sheline YI, Makhoul W, Batzdorf AS, et al. Accelerated Intermittent Theta-Burst Stimulation and Treatment-Refractory Bipolar Depression: A Randomized Clinical Trial. JAMA Psychiatry. 2024;81(9):936–941.

19. Cantone M, Bramanti A, Lanza G, et al. Cortical Plasticity in Depression. ASN Neuro. 2017;9(3):1759091417711512.

20. Lee JC, Corlier J, Wilson AC, et al. Subthreshold stimulation intensity is associated with greater clinical efficacy of intermittent theta-burst stimulation priming for Major Depressive Disorder. Brain Stimulation. 2021;14(4):1015–1021.

21. Solomon EA, Hassan U, Trapp NT, Boes AD, Keller CJ. DLPFC Stimulation Suppresses High-Frequency Neural Activity in the Human sgACC. bioRxiv. Published online April 1, 2025:2025.03.26.645556.

22. Ozdemir RA, Boucher P, Fried PJ, et al. Reproducibility of cortical response modulation induced by intermittent and continuous theta-burst stimulation of the human motor cortex. Brain Stimul. 2021;14(4):949–964.

23. Mittal N, Thakkar B, Hodges CB, et al. Effect of neuroanatomy on corticomotor excitability during and after transcranial magnetic stimulation and intermittent theta burst stimulation. Human Brain Mapping. 2022;43(14):4492–4507.

24. Leodori G, Fabbrini A, De Bartolo MI, et al. Cortical mechanisms underlying variability in intermittent theta-burst stimulation-induced plasticity: A TMS-EEG study. Clinical Neurophysiology. 2021;132(10):2519–2531.

25. Zrenner C, Desideri D, Belardinelli P, Ziemann U. Real-time EEG-defined excitability states determine efficacy of TMS-induced plasticity in human motor cortex. Brain Stimul. 2018;11(2):374–389.

26. Zrenner B, Zrenner C, Gordon PC, et al. Brain oscillation-synchronized stimulation of the left dorsolateral prefrontal cortex in depression using real-time EEG-triggered TMS. Brain Stimulation. 2020;13(1):197–205.

27. Zrenner C, Belardinelli P, Müller-Dahlhaus F, Ziemann U. Closed-Loop Neuroscience and Non-Invasive Brain Stimulation: A Tale of Two Loops. Front Cell Neurosci. 2016;10.

28. Hassan U, Okyere P, Masouleh MA, Zrenner C, Ziemann U, Bergmann TO. Pulsed inhibition of corticospinal excitability by the thalamocortical sleep spindle. Brain Stimul. 2025;18(2):265–275.

29. Stefanou MI, Desideri D, Belardinelli P, Zrenner C, Ziemann U. Phase Synchronicity of μ-Rhythm Determines Efficacy of Interhemispheric Communication Between Human Motor Cortices. J Neurosci. 2018;38(49):10525–10534.

30. Momi D, Ozdemir RA, Tadayon E, et al. Phase-dependent local brain states determine the impact of image-guided transcranial magnetic stimulation on motor network electroencephalographic synchronization. The Journal of Physiology. 2022;600(6):1455–1471.

31. Fujioka T, Trainor LJ, Large EW, Ross B. Internalized Timing of Isochronous Sounds Is Represented in Neuromagnetic Beta Oscillations. J Neurosci. 2012;32(5):1791–1802.

32. Fujioka T, Ross B, Trainor LJ. Beta-Band Oscillations Represent Auditory Beat and Its Metrical Hierarchy in Perception and Imagery. J Neurosci. 2015;35(45):15187–15198.

33. Iversen JR, Repp BH, Patel AD. Top-Down Control of Rhythm Perception Modulates Early Auditory Responses. Ann NY Acad Sci. 2009;1169(1):58–73.

34. Varlet M, Nozaradan S, Trainor L, Keller PE. Dynamic Modulation of Beta Band Cortico-Muscular Coupling Induced by Audio-Visual Rhythms. Cereb Cortex Commun. 2020;1(1):tgaa043.

35. Saleh M, Reimer J, Penn R, Ojakangas CL, Hatsopoulos NG. Fast and slow oscillations in human primary motor cortex predict oncoming behaviorally relevant cues. Neuron. 2010;65(4):461–471.

36. Snyder JS, Large EW. Gamma-band activity reflects the metric structure of rhythmic tone sequences. Brain Res Cogn Brain Res. 2005;24(1):117–126.

37. Comstock DC, Balasubramaniam R. Differential motor system entrainment to auditory and visual rhythms. J Neurophysiol. 2022;128(2):326–335.

38. Ross JM, Comstock DC, Iversen JR, Makeig S, Balasubramaniam R. Cortical mu rhythms during action and passive music listening. J Neurophysiol. 2022;127(1):213–224.

39. Comstock DC, Ross JM, Balasubramaniam R. Modality-specific frequency band activity during neural entrainment to auditory and visual rhythms. Foxe J, ed. Eur J Neurosci. 2021;54(2):4649–4669.

40. Fujioka T, Trainor LJ, Large EW, Ross B. Beta and Gamma Rhythms in Human Auditory Cortex during Musical Beat Processing. Ann NY Acad Sci. 2009;1169(1):89–92.

41. Stupacher J, Hove MJ, Novembre G, Schütz-Bosbach S, Keller PE. Musical groove modulates motor cortex excitability: A TMS investigation. Brain and Cognition. 2013;82(2):127–136.

42. Smayda KE, Cooper SH, Leyden K, Ulaszek J, Ferko N, Dobrin A. Validating the Safe and Effective Use of a Neurorehabilitation System (InTandem) to Improve Walking in the Chronic Stroke Population: Usability Study. JMIR Rehabil Assist Technol. 2023;10:e50438.

43. Awad LN, Jayaraman A, Nolan KJ, et al. Efficacy and safety of using auditory-motor entrainment to improve walking after stroke: a multi-site randomized controlled trial of InTandemTM. Nat Commun. 2024;15(1):1081.

44. Ross JM, Forman L, Gogulski J, et al. Sensory Entrained TMS (seTMS) Enhances Motor Cortex Excitability. Human Brain Mapping. 2025;46(10):e70267.

45. Rossi S, Antal A, Bestmann S, et al. Safety and recommendations for TMS use in healthy subjects and patient populations, with updates on training, ethical and regulatory issues: Expert Guidelines. Clinical Neurophysiology. 2021;132(1):269–306.

46. Rossi S, Hallett M, Rossini PM, Pascual-Leone A. Screening questionnaire before TMS: An update. Clinical Neurophysiology. 2011;122(8):1686.

47. Yeung A, Feldman G, Pedrelli P, et al. The Quick Inventory of Depressive Symptomatology, clinician rated and self-report: a psychometric assessment in Chinese Americans with major depressive disorder. J Nerv Ment Dis. 2012;200(8):712–715.

48. Rush AJ, Trivedi MH, Ibrahim HM, et al. The 16-Item Quick Inventory of Depressive Symptomatology (QIDS), clinician rating (QIDS-C), and self-report (QIDS-SR): a psychometric evaluation in patients with chronic major depression. Biol Psychiatry. 2003;54(5):573–583.

49. Saatlou FH, Rogasch NC, McNair NA, et al. MAGIC: An open-source MATLAB toolbox for external control of transcranial magnetic stimulation devices. Brain Stimulation. 2018;11(5):1189–1191.

50. Hassan U, Pillen S, Zrenner C, Bergmann TO. The Brain Electrophysiological recording & STimulation (BEST) toolbox. Brain Stimul. 2022;15(1):109–115.

51. Rossi S, Hallett M, Rossini PM, Pascual-Leone A, Safety of TMS Consensus Group. Safety, ethical considerations, and application guidelines for the use of transcranial magnetic stimulation in clinical practice and research. Clin Neurophysiol. 2009;120(12):2008–2039.

52. Rossini PM, Barker AT, Berardelli A, et al. Non-invasive electrical and magnetic stimulation of the brain, spinal cord and roots: basic principles and procedures for routine clinical application. Report of an IFCN committee. Electroencephalography and Clinical Neurophysiology. 1994;91(2):79–92.

53. Rossini PM, Burke D, Chen R, et al. Non-invasive electrical and magnetic stimulation of the brain, spinal cord, roots and peripheral nerves: Basic principles and procedures for routine clinical and research application. An updated report from an I.F.C.N. Committee. Clinical Neurophysiology. 2015;126(6):1071–1107.

54. Stokes MG, Chambers CD, Gould IC, et al. Simple Metric For Scaling Motor Threshold Based on Scalp-Cortex Distance: Application to Studies Using Transcranial Magnetic Stimulation. Journal of Neurophysiology. 2005;94(6):4520–4527.

55. Pridmore S, Fernandes Filho JA, Nahas Z, Liberatos C, George MS. Motor threshold in transcranial magnetic stimulation: a comparison of a neurophysiological method and a visualization of movement method. J ECT. 1998;14(1):25–27.

56. Wu SW, Shahana N, Huddleston DA, Gilbert DL. Effects of 30Hz θ burst transcranial magnetic stimulation on the primary motor cortex. J Neurosci Methods. 2012;208(2):161–164.

57. Todd G, Flavel SC, Ridding MC. Priming theta-burst repetitive transcranial magnetic stimulation with low- and high-frequency stimulation. Exp Brain Res. 2009;195(2):307–315.

58. Gedankien T, Fried PJ, Pascual-Leone A, Shafi MM. Intermittent theta-burst stimulation induces correlated changes in cortical and corticospinal excitability in healthy older subjects. Clin Neurophysiol. 2017;128(12):2419–2427.

59. Repp BH. Rate Limits of On-Beat and Off-Beat Tapping With Simple Auditory Rhythms. Music Perception. 2005;23(2):165–188.

60. Repp BH, Su YH. Sensorimotor synchronization: A review of recent research (2006–2012). Psychon Bull Rev. 2013;20(3):403–452.

61. Ross JM, Iversen JR, Balasubramaniam R. Motor simulation theories of musical beat perception. Neurocase. 2016;22(6):558–565.

62. Janata P, Tomic ST, Haberman JM. Sensorimotor coupling in music and the psychology of the groove. J Exp Psychol. 2012;141(1):54–75.

63. Stupacher J, Hove MJ, Janata P. Audio Features Underlying Perceived Groove and Sensorimotor Synchronization in Music. Music Percept. 2016;33(5):571–589.

64. Madison G. Experiencing Groove Induced by Music: Consistency and Phenomenology. Music Perception. 2006;24(2):201–208.

65. Nombela C, Hughes LE, Owen AM, Grahn JA. Into the groove: Can rhythm influence Parkinson’s disease? Neuroscience & Biobehavioral Reviews. 2013;37(10):2564–2570.

66. Ross JM, Warlaumont AS, Abney DH, Rigoli LM, Balasubramaniam R. Influence of musical groove on postural sway. J Exp Psychol Hum Percept Perform. 2016;42(3):308–319.

67. Schilberg L, Ten Oever S, Schuhmann T, Sack AT. Phase and power modulations on the amplitude of TMS-induced motor evoked potentials. PLoS One. 2021;16(9):e0255815.

68. Guidali G, Zazio A, Lucarelli D, et al. Effects of transcranial magnetic stimulation (TMS) current direction and pulse waveform on cortico-cortical connectivity: A registered report TMS-EEG study. Eur J of Neuroscience. 2023;58(8):3785–3809.

69. Spampinato DA, Ibanez J, Rocchi L, Rothwell J. Motor potentials evoked by transcranial magnetic stimulation: interpreting a simple measure of a complex system. J Physiol. 2023;601(14):2827–2851.

70. Dunn OJ. Multiple Comparisons among Means. Journal of the American Statistical Association. 1961;56(293):52–64.

71. Chung SW, Hill AT, Rogasch NC, Hoy KE, Fitzgerald PB. Use of theta-burst stimulation in changing excitability of motor cortex: A systematic review and meta-analysis. Neuroscience & Biobehavioral Reviews. 2016;63:43–64.

72. Bates D, Mächler M, Bolker B, Walker S. Fitting Linear Mixed-Effects Models Using lme4. J Stat Soft. 2015;67(1).

73. Luke SG. Evaluating significance in linear mixed-effects models in R. Behav Res. 2017;49(4):1494–1502.

74. Little RJA, Rubin DB. Statistical Analysis with Missing Data. Second edition. Wiley-Interscience; 2002.

75. Lenth RV. emmeans: Estimated Marginal Means, aka Least-Squares Means. Published online October 20, 2017:1.10.6.

76. Pellicciari MC, Miniussi C, Ferrari C, Koch G, Bortoletto M. Ongoing cumulative effects of single TMS pulses on corticospinal excitability: An intra- and inter-block investigation. Clin Neurophysiol. 2016;127(1):621–628.

77. Veniero D, Bortoletto M, Miniussi C. TMS-EEG co-registration: On TMS-induced artifact. Clinical Neurophysiology. 2009;120(7):1392–1399.

78. Ross JM, Santarnecchi E, Lian SJ, et al. Neurophysiologic predictors of individual risk for post-operative delirium after elective surgery. J American Geriatrics Society. 2023;71(1):235–244.

79. Liu C, Wechsler H. Comparative Assessment of Independent Component Analysis (ICA) for Face Recognition. Published online 1999:6.

80. Draper BA, Baek K, Bartlett MS, Beveridge JR. Recognizing faces with PCA and ICA. Computer Vision and Image Understanding. 2003;91(1-2):115–137.

81. Artoni F, Delorme A, Makeig S. Applying dimension reduction to EEG data by Principal Component Analysis reduces the quality of its subsequent Independent Component decomposition. Neuroimage. 2018;175:176–187.

82. Hyvarinen A. Fast and robust fixed-point algorithms for independent component analysis. IEEE Trans Neural Netw. 1999;10(3):626–634.

83. Winkler I, Haufe S, Tangermann M. Automatic classification of artifactual ICA-components for artifact removal in EEG signals. Behav Brain Funct. 2011;7:30.

84. Winkler I, Brandl S, Horn F, Waldburger E, Allefeld C, Tangermann M. Robust artifactual independent component classification for BCI practitioners. J Neural Eng. 2014;11(3):035013.

85. Rogasch NC, Sullivan C, Thomson RH, et al. Analysing concurrent transcranial magnetic stimulation and electroencephalographic data: A review and introduction to the open-source TESA software. NeuroImage. 2017;147:934–951.

86. Mutanen TP, Biabani M, Sarvas J, Ilmoniemi RJ, Rogasch NC. Source-based artifact-rejection techniques available in TESA, an open-source TMS–EEG toolbox. Brain Stimulation. 2020;13(5):1349–1351.

87. Pion-Tonachini L, Kreutz-Delgado K, Makeig S. ICLabel: An automated electroencephalographic independent component classifier, dataset, and website. Neuroimage. 2019;198:181–197.

88. Pineda JA. The functional significance of mu rhythms: Translating “seeing” and “hearing” into “doing.” Brain Research Reviews. 2005;50(1):57–68.

89. Singh F, Pineda J, Cadenhead KS. Association of impaired EEG mu wave suppression, negative symptoms and social functioning in biological motion processing in first episode of psychosis. Schizophr Res. 2011;130(1-3):182–186.

90. DiGirolamo MA, Simon JC, Hubley KM, Kopulsky A, Gutsell JN. Clarifying the relationship between trait empathy and action-based resonance indexed by EEG mu-rhythm suppression. Neuropsychologia. 2019;133:107172.

91. Pfurtscheller G, Stancák A, Neuper C. Event-related synchronization (ERS) in the alpha band--an electrophysiological correlate of cortical idling: a review. Int J Psychophysiol. 1996;24(1-2):39–46.

92. Pfurtscheller G, Neuper C, Andrew C, Edlinger G. Foot and hand area mu rhythms. Int J Psychophysiol. 1997;26(1-3):121–135.

93. Delorme A, Makeig S. EEGLAB: an open source toolbox for analysis of single-trial EEG dynamics including independent component analysis. Journal of Neuroscience Methods. 2004;134(1):9–21.

94. Makeig S. Auditory event-related dynamics of the EEG spectrum and effects of exposure to tones. Electroencephalography and Clinical Neurophysiology. 1993;86(4):283–293.

95. Grandchamp R, Delorme A. Single-Trial Normalization for Event-Related Spectral Decomposition Reduces Sensitivity to Noisy Trials. Front Psychology. 2011;2.

96. Wisniewski MG, Joyner CN, Zakrzewski AC, Makeig S. Finding tau rhythms in EEG: An independent component analysis approach. Hum Brain Mapp. 2024;45(2):e26572.

97. Pfurtscheller G, Lopes da Silva FH. Event-related EEG/MEG synchronization and desynchronization: basic principles. Clin Neurophysiol. 1999;110(11):1842–1857.

98. Hamada M, Murase N, Hasan A, Balaratnam M, Rothwell JC. The role of interneuron networks in driving human motor cortical plasticity. Cereb Cortex. 2013;23(7):1593–1605.

99. Balasubramaniam R, Haegens S, Jazayeri M, Merchant H, Sternad D, Song JH. Neural Encoding and Representation of Time for Sensorimotor Control and Learning. J Neurosci. 2021;41(5):866–872.

100. Gordon CL, Cobb PR, Balasubramaniam R. Recruitment of the motor system during music listening: An ALE meta-analysis of fMRI data. Grahn JA, ed. PLoS ONE. 2018;13(11):e0207213.

101. Repp BH. Sensorimotor synchronization: A review of the tapping literature. Psychon Bull Rev. 2005;12(6):969–992.

102. Ross JM, Balasubramaniam R. Time Perception for Musical Rhythms: Sensorimotor Perspectives on Entrainment, Simulation, and Prediction. Front Integr Neurosci. 2022;16:916220.

103. Ross JM, Balasubramaniam R. Physical and neural entrainment to rhythm: human sensorimotor coordination across tasks and effector systems. Front Hum Neurosci. 2014;8.

104. Patel AD, Iversen JR. The evolutionary neuroscience of musical beat perception: the Action Simulation for Auditory Prediction (ASAP) hypothesis. Front Syst Neurosci. 2014;8.

105. Grahn JA, Brett M. Impairment of beat-based rhythm discrimination in Parkinson’s disease. Cortex. 2009;45(1):54–61.

106. Grube M, Cooper FE, Chinnery PF, Griffiths TD. Dissociation of duration-based and beat-based auditory timing in cerebellar degeneration. Proc Natl Acad Sci. 2010;107(25):11597–11601.

107. Kotz SA, Brown RM, Schwartze M. Cortico-striatal circuits and the timing of action and perception. Curr Opin Behav Sci. 2016;8:42–45.

108. Grahn JA, Rowe JB. Finding and Feeling the Musical Beat: Striatal Dissociations between Detection and Prediction of Regularity. Cereb Cortex. 2013;23(4):913–921.

109. Ross JM, Iversen JR, Balasubramaniam R. The Role of Posterior Parietal Cortex in Beat-based Timing Perception: A Continuous Theta Burst Stimulation Study. J Cogn Neurosci. 2018;30(5):634–643.

110. Ross J, Iversen J, Balasubramaniam R. Dorsal Premotor Contributions to Auditory Rhythm Perception: Causal Transcranial Magnetic Stimulation Studies of Interval, Tempo, and Phase. bioRxiv. Published online July 13, 2018.

111. Grube M, Lee KH, Griffiths TD, Barker AT, Woodruff PW. Transcranial magnetic theta-burst stimulation of the human cerebellum distinguishes absolute, duration-based from relative, beat-based perception of subsecond time intervals. Front Psychol. 2010;1:171.

112. Pollok B, Rothkegel H, Schnitzler A, Paulus W, Lang N. The effect of rTMS over left and right dorsolateral premotor cortex on movement timing of either hand. Eur J Neurosci. 2008;27(3):757–764.

113. Wang JB, Hassan U, Bruss JE, et al. Effects of transcranial magnetic stimulation on the human brain recorded with intracranial electrocorticography. Mol Psychiatry. 2024;29(5):1228–1240.

114. Hassan U. Intracranial Recording During TMS: A Practical Guide. Preprint posted online February 21, 2025.

115. Spiech C, Danielsen A, Laeng B, Endestad T. Oscillatory attention in groove. Cortex. 2024;174:137–148.

116. Mathias B, Zamm A, Gianferrara PG, Ross B, Palmer C. Rhythm Complexity Modulates Behavioral and Neural Dynamics During Auditory–Motor Synchronization. Journal of Cognitive Neuroscience. 2020;32(10):1864–1880.

117. Zalta A, Large EW, Schön D, Morillon B. Neural dynamics of predictive timing and motor engagement in music listening. Sci Adv. 2024;10(10):eadi2525.

118. Keller CJ, Ross JM. Systems and methods for sensory entrained transcranial magnetic stimulation. Published online August 29, 2024:US-20240285964-A1.

