## Supplement for "Sensory Entrained TMS (seTMS) enhances motor cortex plasticity"

***Table S1. Demographics.*** *n*=20

| | |
|---|---|
| Age, mean years (SD) | 41.0 (14.5) |
| Sex | |
| Female, *n* (%) | 10 (50.0) |
| Male, *n* (%) | 10 (50.0) |
| Other or prefer not to state, *n* (%) | 0 (0.0) |
| Handedness | |
| Left hand dominant, *n* (%) | 1 (5.0) |
| Right hand dominant, *n* (%) | 19 (95.0) |
| Ambidextrous, *n* (%) | 0 (0.0) |
| Education | |
| GED or High School Diploma, *n* (%) | 1 (5.0) |
| Some college, no degree, *n* (%) | 0 (0.0) |
| Two year degree, *n* (%) | 2 (10.0) |
| Four year degree, *n* (%) | 10 (50.0) |
| Post graduate degree, *n* (%) | 7 (35.0) |
| Employment | |
| Part-time, *n* (%) | 6 (22.2) |
| Full-time, *n* (%) | 7 (25.9) |
| Unemployed, *n* (%) | 4 (14.8) |
| Retired, *n* (%) | 1 (3.7) |
| Part-time student, *n* (%) | 0 (0.0) |
| Full-time student, *n* (%) | 2 (7.4) |
| Race | |
| White, *n* (%) | 10 (37.0) |
| Black or African American, *n* (%) | 3 (11.1) |
| American Indian or Alaska Native, *n* (%) | 0 (0.0) |
| Asian, *n* (%) | 6 (22.2) |
| Native Hawaiian or Other Pacific Islander, *n* (%) | 0 (0.0) |
| Some other race or prefer not to state, *n* (%) | 1 (3.7) |

***Table S2. Musical stimuli.***

| name | *artist* | *Groove rating** |
|---|---|---|
| Music | Leela James | 101.1 |
| Outa-Space | Billy Preston | 90.9 |
| Baby It's You | JoJo | 79.7 |

**Groove ratings are from Janata et al. 2012 and are on a scale from 0 to 127*

***Table S3. Comparison of MEP size to baseline.*** *Paired t-tests with baseline at each time point.*

| protocol | time | *n* | *df* | *t* | *h** | *p* | Cohen's *d* | | mean | std. dev. | std. err | 95% CI |
|---|---|---|---|---|---|---|---|---|---|---|---|---|
| **se-iTBS** | | | | | | | | base | 1.07 | 0.04 | 0.009 | [1.05 1.09] |
| | **t0** | **20** | **19** | **-6.82** | **1** | **0.000002** | **-1.52** | | 1.07 | 0.04 | 0.009 | [1.39 1.70] |
| | **t15** | **20** | **19** | **-4.29** | **1** | **0.0004** | **-0.96** | | 1.07 | 0.04 | 0.009 | [1.24 1.60] |
| | **t30** | **20** | **19** | **-4.23** | **1** | **0.0004** | **-0.95** | | 1.07 | 0.04 | 0.009 | [1.24 1.60] |
| | t45 | $10^+$ | 9 | -1.54 | 0 | 0.16 | | | 1.07 | 0.04 | 0.009 | [1.07 1.26] |
| **iTBS** | | | | | | | | base | 1.07 | 0.06 | 0.01 | [1.04 1.10] |
| | **t0** | **20** | **19** | **-4.18** | **1** | **0.0005** | **-0.94** | | 1.07 | 0.06 | 0.01 | [1.16 1.36] |
| | t15 | 20 | 19 | -2.47 | 0 | 0.02 | | | 1.07 | 0.06 | 0.01 | [1.08 1.31] |
| | t30 | 20 | 19 | -1.78 | 0 | 0.09 | | | 1.07 | 0.06 | 0.01 | [1.05 1.28] |
| | t45 | $10^+$ | 9 | 0.46 | 0 | 0.65 | | | 1.07 | 0.06 | 0.01 | [0.93 1.17] |

**When corrected for 8 tests using the Bonferroni method, $\alpha = 0.006$ (=0.05 / 8)*

+Very small sample size

***Table S4. Average percent increase in MEPs from baseline.*** *Post-iTBS compared with pre-iTBS.*

| protocol | *0 min* | *15 min* | *30 min* | *45 min* |
|---|---|---|---|---|
| se-iTBS | 54.58 | 42.17 | 36.67 | 16.79 |
| iTBS | 25.97 | 19.34 | 16.66 | 4.87 |

***Table S5. Average percent increase in MEP modulation when using se-iTBS.*** *Compared with standard iTBS.*

| time | *% increase* |
|---|---|
| t0 | 22.71 |
| t15 | 19.13 |
| t30 | 17.15 |
| t45 | 11.37 |

***Table S6. Nonparametric comparison of MEP modulation.*** *Wilcoxon signed rank tests at each time point.*

| time | *n* | *df* | *h* | *p* |
|---|---|---|---|---|
| **t0** | **20** | **19** | **1** | **0.0009** |
| **t15** | **20** | **19** | **1** | **0.0005** |
| **t30** | **20** | **19** | **1** | **0.006** |
| **t45** | **$10^+$** | **9** | **1** | **0.04** |

***Table S7. Comparison of baseline MEPs.*** *Paired t-test.*

| time | *n* | *df* | *t* | *h* | *p* | Cohen's *d* | | mean | std. dev. | std. err | 95% CI |
|---|---|---|---|---|---|---|---|---|---|---|---|
| base | 20 | 19 | -0.26 | 0 | 0.80 | -0.06 | | | | | |
| | | | | | | | seTMS | 1.07 | 0.04 | 0.009 | [ 1.05 1.09] |
| | | | | | | | TMS | 1.07 | 0.06 | 0.012 | [ 1.04 1.10] |

***Table S8. Non-parametric comparison of baseline MEPs.*** *Wilcoxon signed rank test.*

| time | *n* | *df* | *h* | *p* |
|---|---|---|---|---|
| base | 20 | 19 | 0 | 0.82 |

***Table S9. Comparison of ITC trough latencies.*** *Paired t-tests.*

| Grouping | *n* | *df* | *t* | *h* | *p* | Cohen's *d* | | mean | std. dev. | std. err | 95% CI |
|---|---|---|---|---|---|---|---|---|---|---|---|
| by protocol | 20 | 19 | -1.39 | 0 | 0.18 | -0.31 | | | | | |
| | | | | | | | pre-se-iTBS | -172.05 | 44.44 | 9.94 | [-192.85 -151.25] |
| | | | | | | | pre-iTBS | -151.70 | 42.40 | 9.48 | [-171.55 -131.85] |
| by order | 20 | 19 | -0.36 | 0 | 0.72 | -0.08 | | | | | |
| | | | | | | | first | -164.60 | 43.32 | 9.69 | [-184.88 -144.32] |
| | | | | | | | second | -159.15 | 45.81 | 10.24 | [-180.59 -137.71] |

***Table S10. Nonparametric comparison of ITC trough latencies.*** *Wilcoxon signed rank tests.*

| Grouping | *n* | *df* | *h* | *p* |
|---|---|---|---|---|
| by protocol | 20 | 19 | 0 | 0.17 |
| by order | 20 | 19 | 0 | 0.70 |

***Table S11. Exploratory regressions**. Individual ITC metrics (trough timing, distance from -200 ms, ITC value at trough, y-intercept and slope of the fitted ITC line) do not predict MEP outcomes (plasticity at 0, 15, and 30 min; max and mean plasticity from se-iTBS; se-iTBS gain over standard iTBS). No significant relationships emerged (all p > 0.05, uncorrected). This null result likely reflects limited variability in trough-targeting quality across participants. If the group-based -200 ms timing already approximates individual troughs reasonably well, the predictor range would be compressed, reducing power to detect individual-difference effects. Additionally, ITC metrics derived from EEG carry inherent measurement noise, and even if reliably measured, the relationship between ITC metrics and MEP outcomes may not follow a simple linear pattern.*

| Predictor | Outcome | $R^2$ | $F$ | $p$ |
|---|---|---|---|---|
| ITC trough time | MEP plasticity at 0min | 0.0379 | 0.631 | 0.439 |
| ITC trough time | MEP plasticity at 15min | 0.0773 | 1.34 | 0.264 |
| ITC trough time | MEP plasticity at 30min | 0.196 | 3.90 | 0.0657 |
| ITC trough time | max plasticity from seTMS | 0.0876 | 1.54 | 0.233 |
| ITC trough time | mean plasticity from seTMS | 0.127 | 2.33 | 0.146 |
| ITC trough time | max seTMS gain over standard TMS | 0.00199 | 0.0319 | 0.860 |
| ITC trough time | avg seTMS gain over standard TMS | 0.000828 | 0.0133 | 0.910 |
| ITC trough distance from -200ms | MEP plasticity at 0min | 0.0484 | 0.814 | 0.380 |
| ITC trough distance from -200ms | MEP plasticity at 15min | 0.141 | 2.64 | 0.124 |
| ITC trough distance from -200ms | MEP plasticity at 30min | 0.212 | 4.31 | 0.0544 |
| ITC trough distance from -200ms | max plasticity from seTMS | 0.158 | 3.01 | 0.102 |
| ITC trough distance from -200ms | mean plasticity from seTMS | 0.173 | 3.34 | 0.0863 |
| ITC trough distance from -200ms | max seTMS gain over standard TMS | 0.0437 | 0.731 | 0.405 |
| ITC trough distance from -200ms | avg seTMS gain over standard TMS | 0.0403 | 0.672 | 0.424 |
| ITC value at trough | MEP plasticity at 0min | 0.0541 | 0.916 | 0.353 |
| ITC value at trough | MEP plasticity at 15min | 0.00337 | 0.0541 | 0.819 |
| ITC value at trough | MEP plasticity at 30min | 0.0174 | 0.283 | 0.602 |
| y intercept of fitted line | MEP plasticity at 0min | 0.00134 | 0.0214 | 0.886 |
| y intercept of fitted line | MEP plasticity at 15min | 0.000818 | 0.0131 | 0.910 |
| y intercept of fitted line | MEP plasticity at 30min | 0.0000143 | 0.000229 | 0.988 |
| slope of fitted line | MEP plasticity at 0min | 0.0521 | 0.880 | 0.362 |
| slope of fitted line | MEP plasticity at 15min | 0.0140 | 0.227 | 0.640 |
| slope of fitted line | MEP plasticity at 30min | 0.102 | 1.81 | 0.197 |

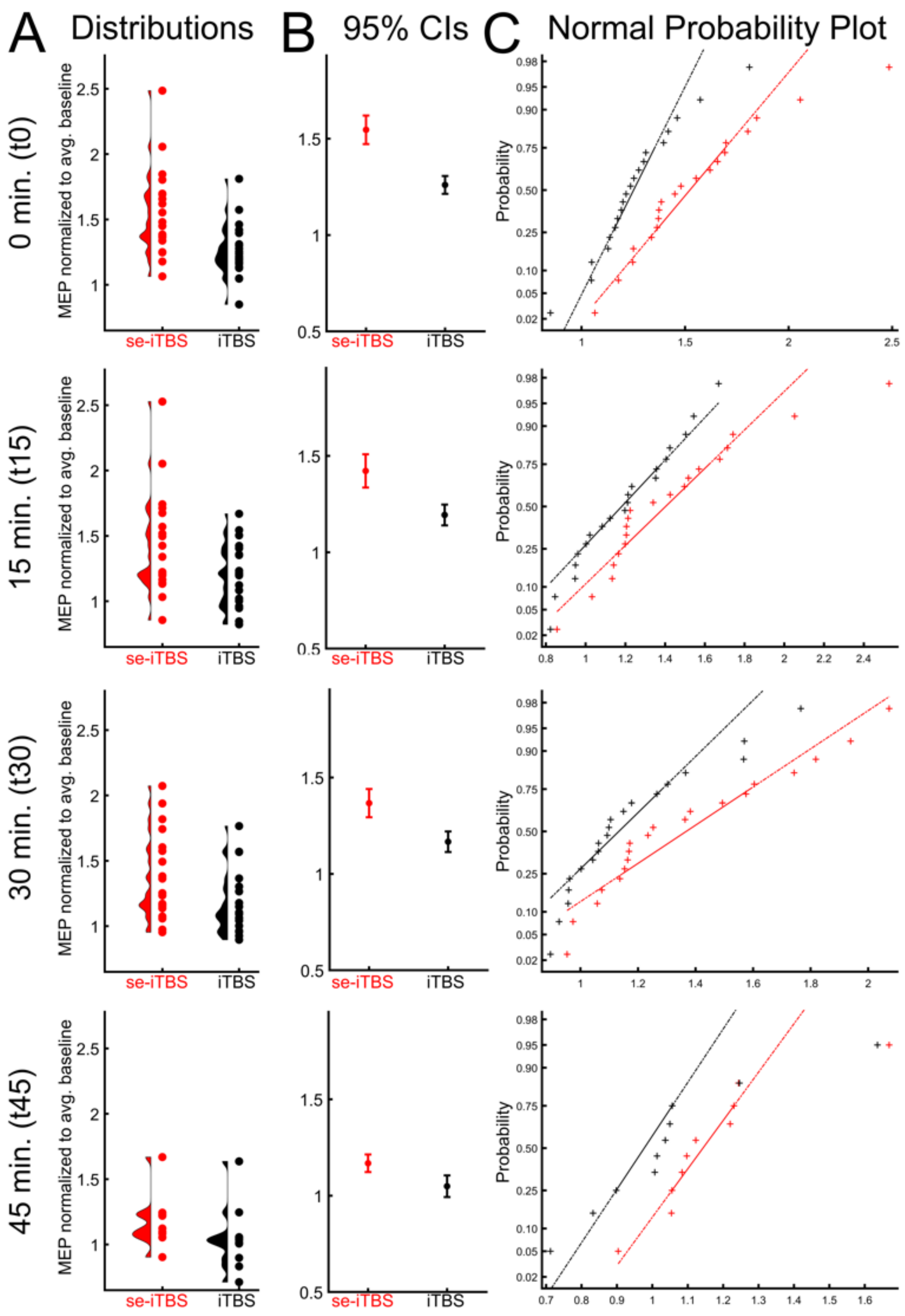


**Figure S1. Review of statistical assumptions.** MEP amplitudes used for paired samples *t*-tests comparing se-iTBS to iTBS within each time point: t0, t15, t30 and t45. MEP amplitudes are normalized to average baseline amplitude. A) Data distributions. B) 95% confidence intervals. C) Normal probability plots showing our data on the x-axis to the normal distribution on the y-axis. A straight line suggests normality.

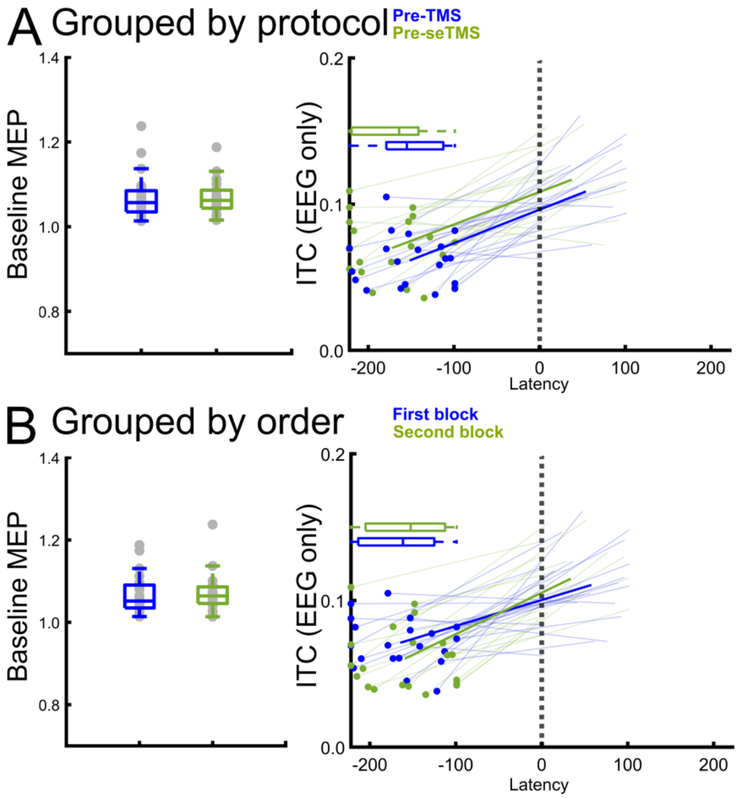


**Figure S2. No differences between baseline MEPs or EEG latencies from the beat time.** A) Grouped by protocol. Baseline normalized MEP amplitudes (left) and ITC trough latencies from EEG recordings (right) confirmed no differences prior to each protocol. B) Grouped by order. Baseline normalized MEP amplitudes and ITC trough latencies from EEG recordings confirmed no differences when grouped by the order they were recorded in the session. Paired samples *t*-tests confirmed that neither baseline MEPs nor ITC latencies were significantly different from each other (Tables S6-9)

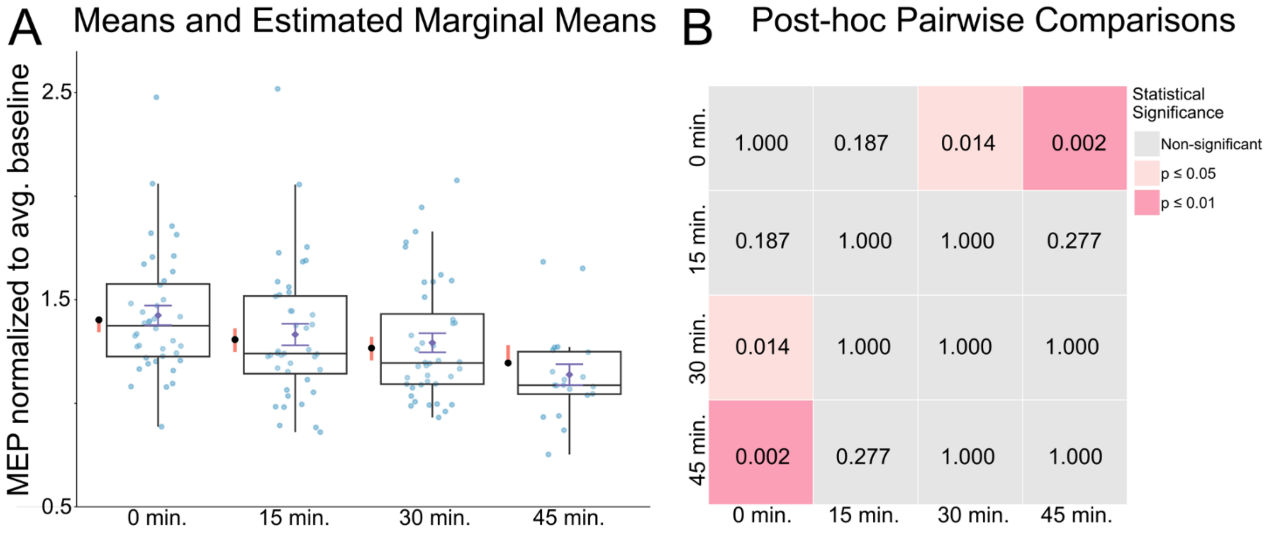


**Figure S3. Linear mixed-effect model demonstrating modulation effect.** The model revealed significant effects of protocol ($F(1,113) = 44.95$, $p<.0001$) and of time point ($F(3,113) = 5.70$, $p=.001$) but no protocol by time point interaction ($F(3,113) = 0.87$, $p=.46$). A) Normalized MEPs for each time point (blue) with box and whisker plots showing the means and standard error, and estimated marginal means are to the left (black) with lower and upper comparison limits (red). B) Post-hoc pairwise comparisons with Bonferroni correction indicated significant differences between time points 0 min and 30 min ($t(113) = 3.12$, $p = 0.01$), and between time points 0 min and 45 min ($t(113) = 3.72$, $p = 0.002$). These post-hoc results suggest that the MEP modulation effects reduced with time after the protocol and, because we did not find a protocol by time point interaction, we can interpret that this did not depend on which protocol was used. See Table 2 for test statistics for all post-hoc comparisons.

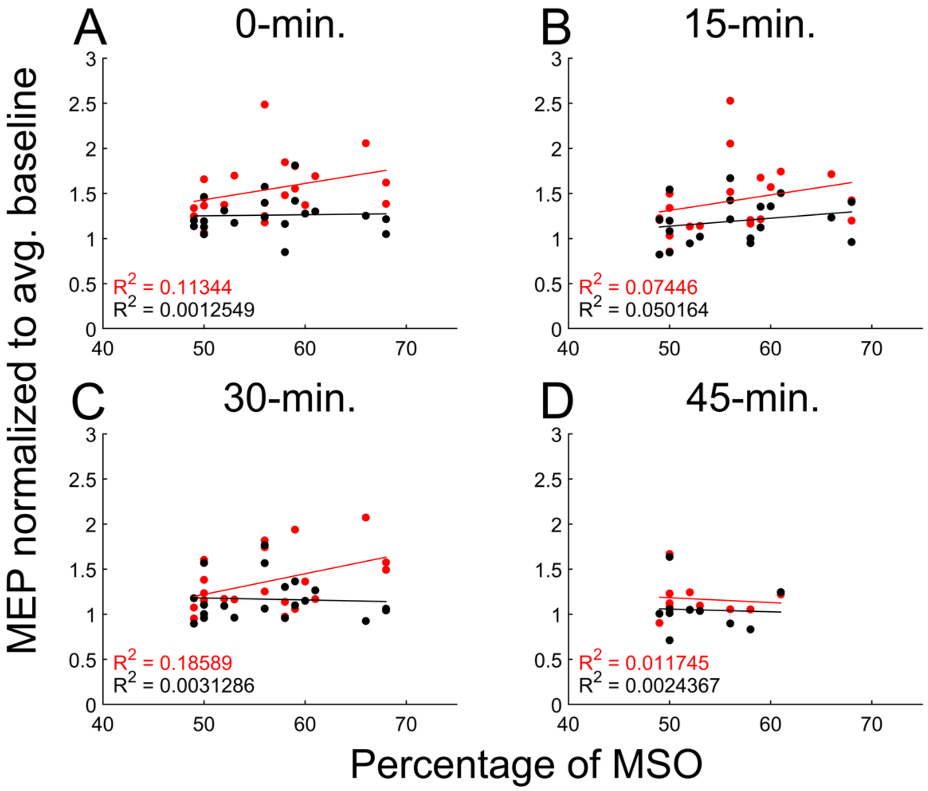


**Figure S4. MEP modulation as a function of maximum stimulator output.** The iTBS protocols were administered at 90% rMT for each participant. Simple linear regressions were used to check for systematic changes in MEP modulation with iTBS intensity when taken as a percentage of maximum stimulator output. No significant relationships were found, at 0-min (se-iTBS $R^2$ = 0.11, $F(2,18)$ = 1.15, $p$ = 0.34; iTBS $R^2$ = 0.0012, $F(2,18)$ = 0.011, $p$ = 0.99), 15-min (se-iTBS $R^2$ = 0.074, $F(2,18)$ = 0.72, $p$ = 0.50; iTBS $R^2$ = 0.050, $F(2,18)$ = 0.48, $p$ = 0.63), 30-min (se-iTBS $R^2$ = 0.18, $F(2,18)$ = 2.06, $p$ = 0.16; iTBS $R^2$ = 0.0031, $F(2,18)$ = 0.028, $p$ = 0.97), or 45-min (se-iTBS $R^2$ = 0.012, $F(2,8)$ = 0.048, $p$ = 0.95; iTBS $R^2$ = 0.0024, $F(2,8)$ = 0.0098, $p$ = 0.99).

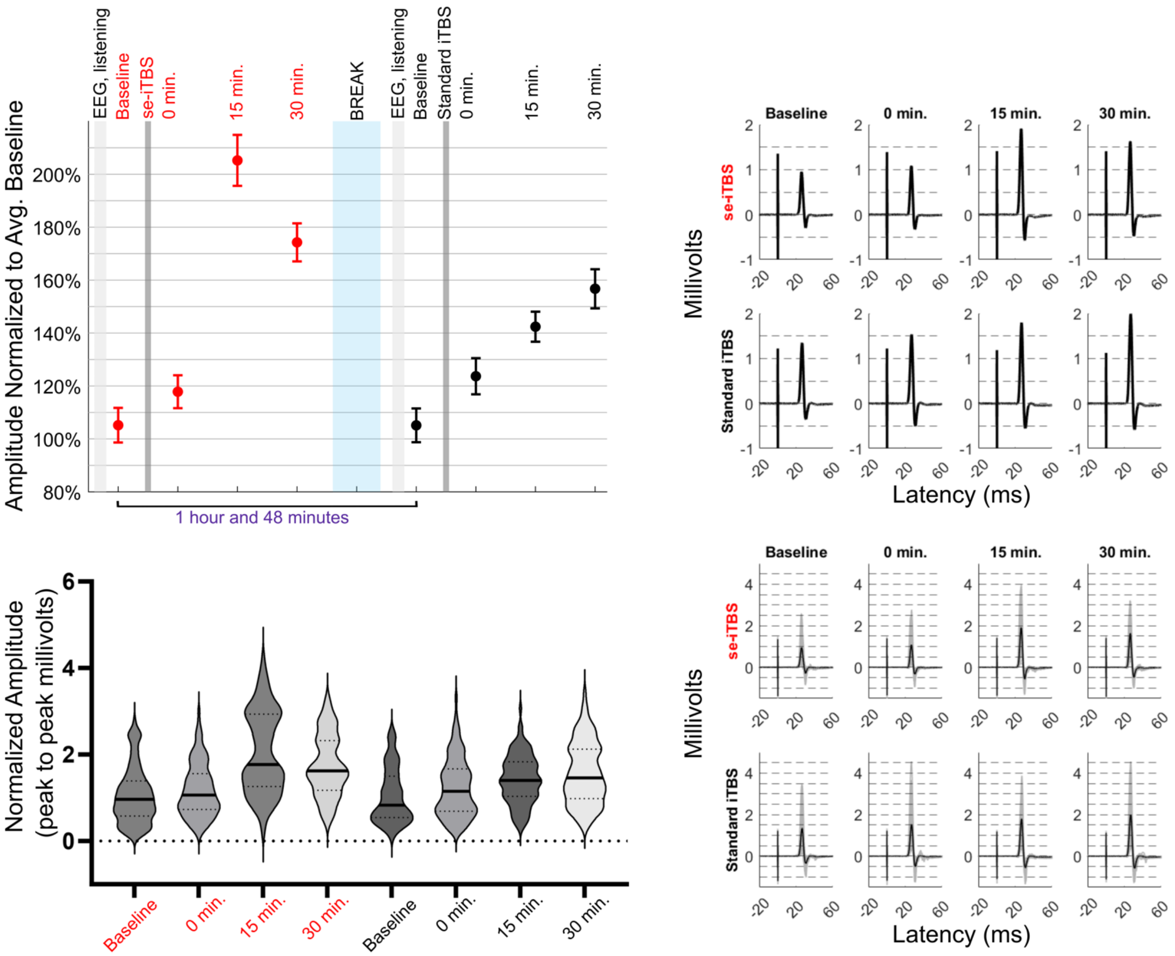


**Figure S5. Participant 1 individual data.**

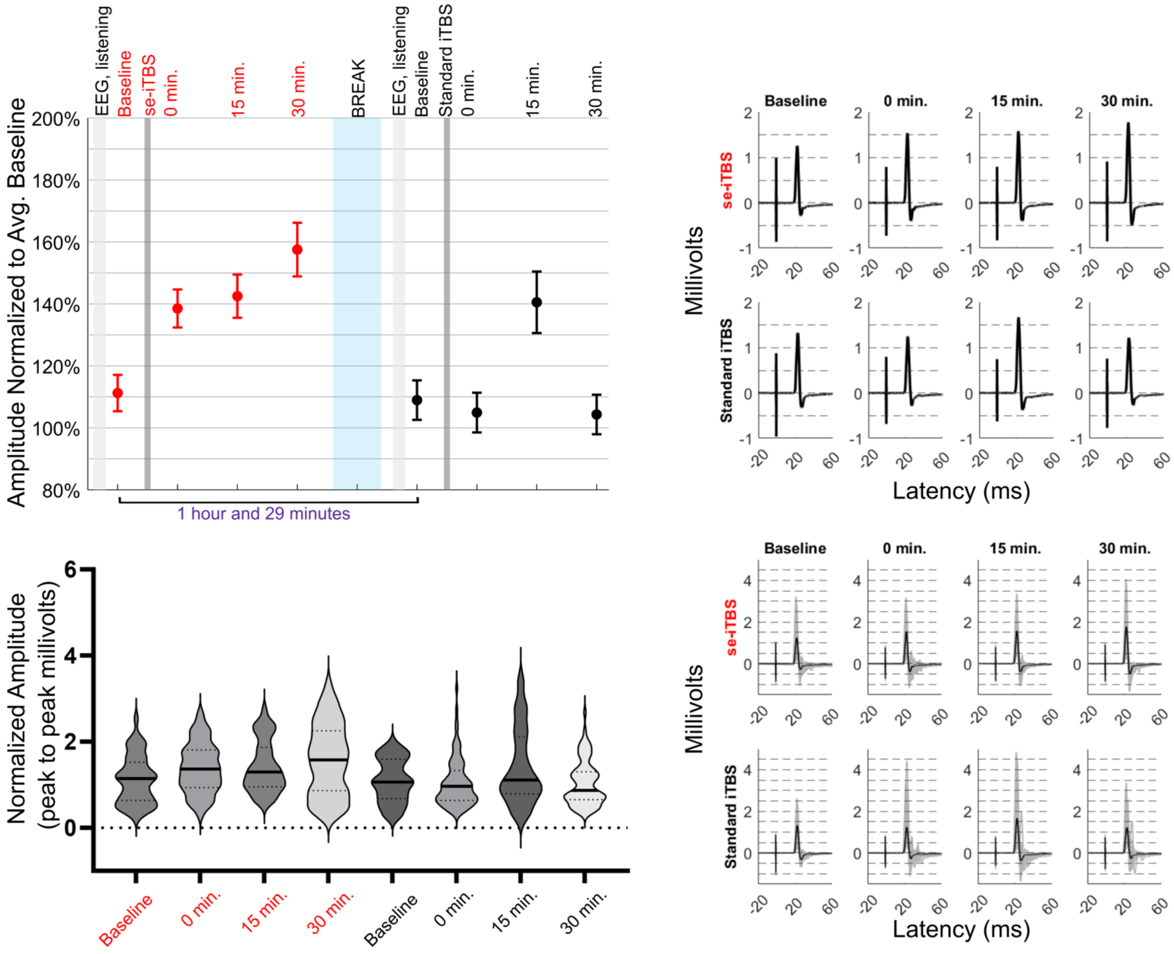


**Figure S6. Participant 2 individual data**

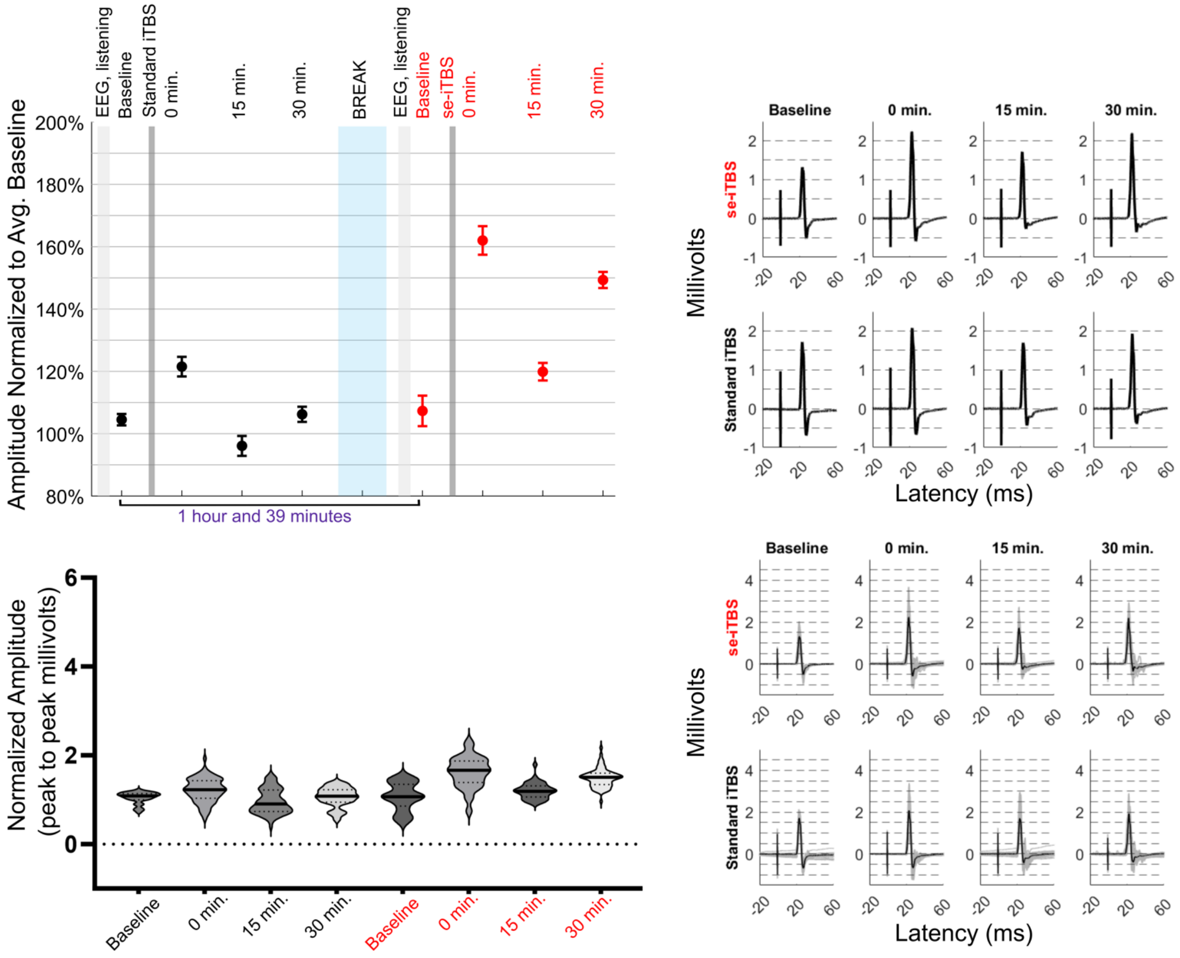


**Figure S7. Participant 3 individual data**

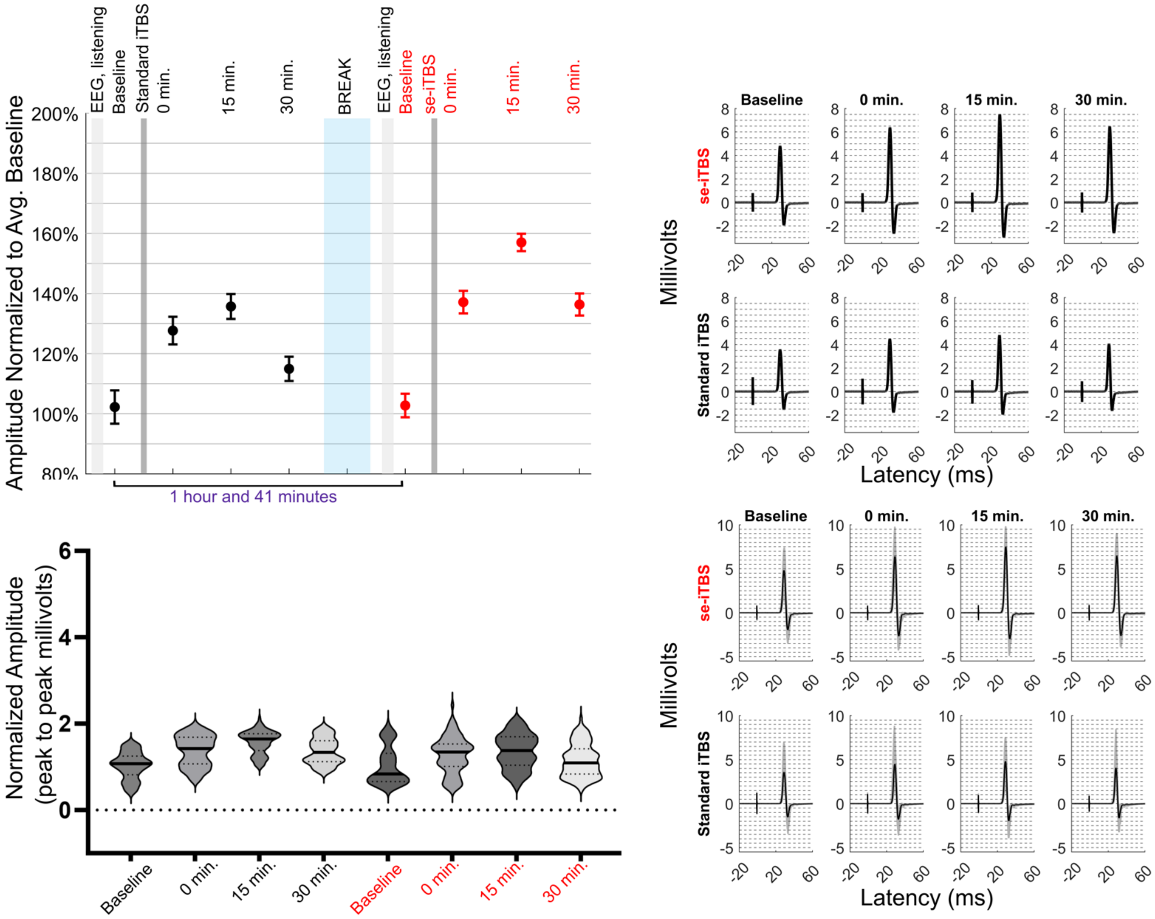


**Figure S8. Participant 4 individual data**

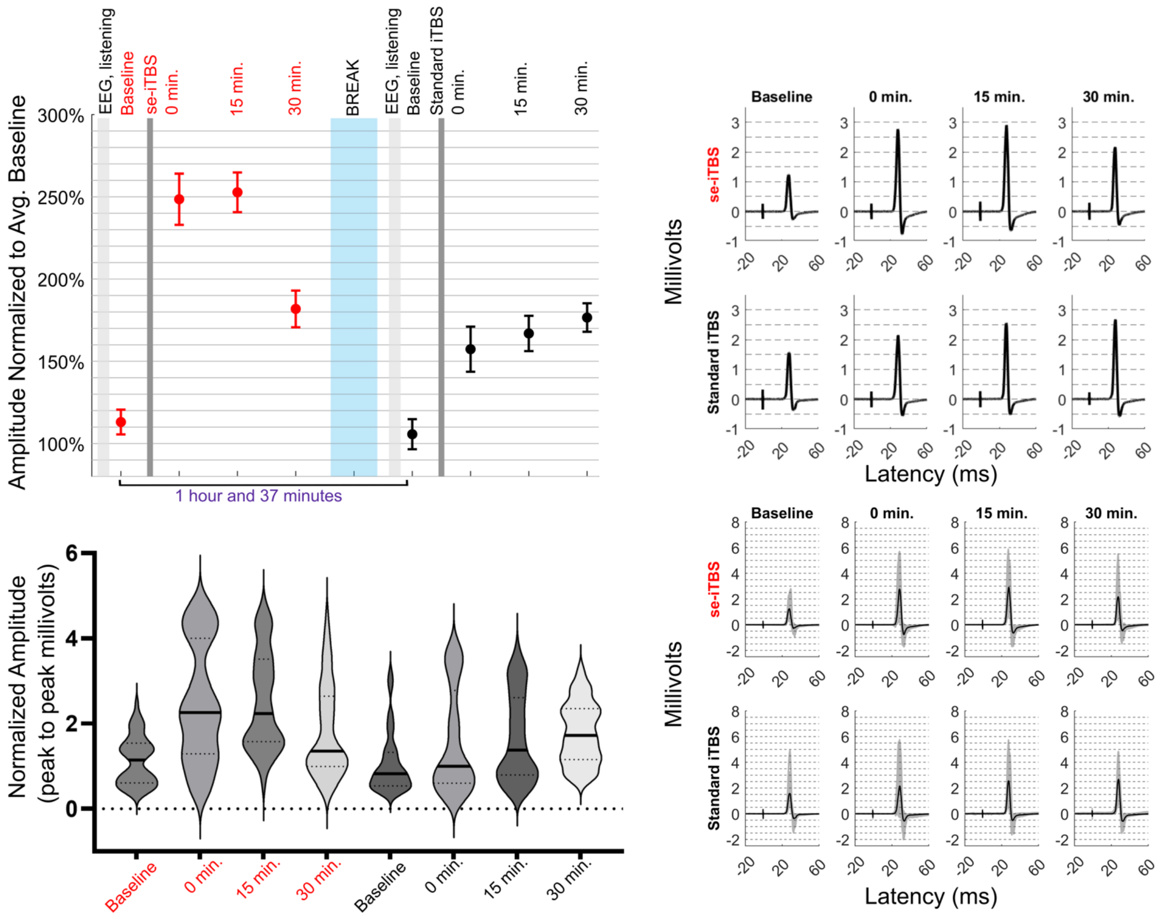


**Figure S9. Participant 5 individual data**

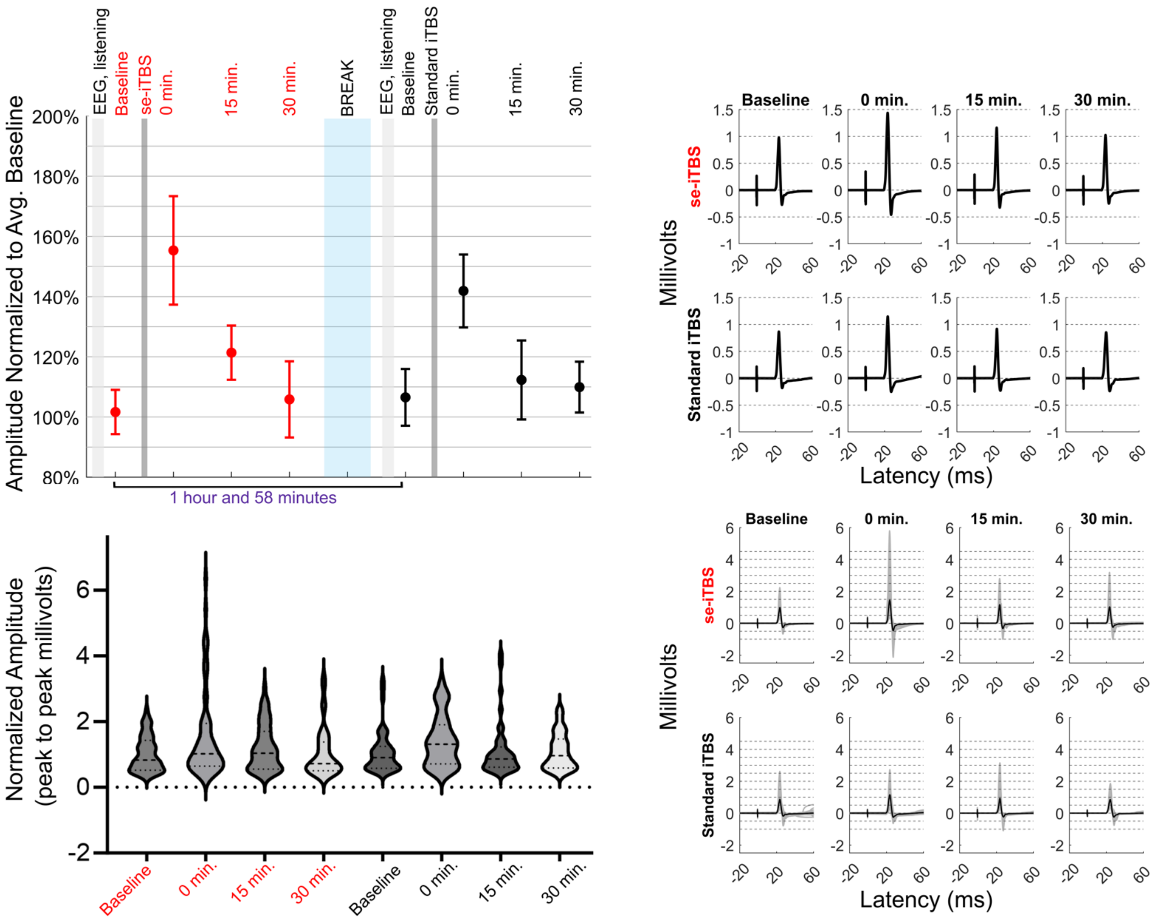


**Figure S10. Participant 6 individual data**

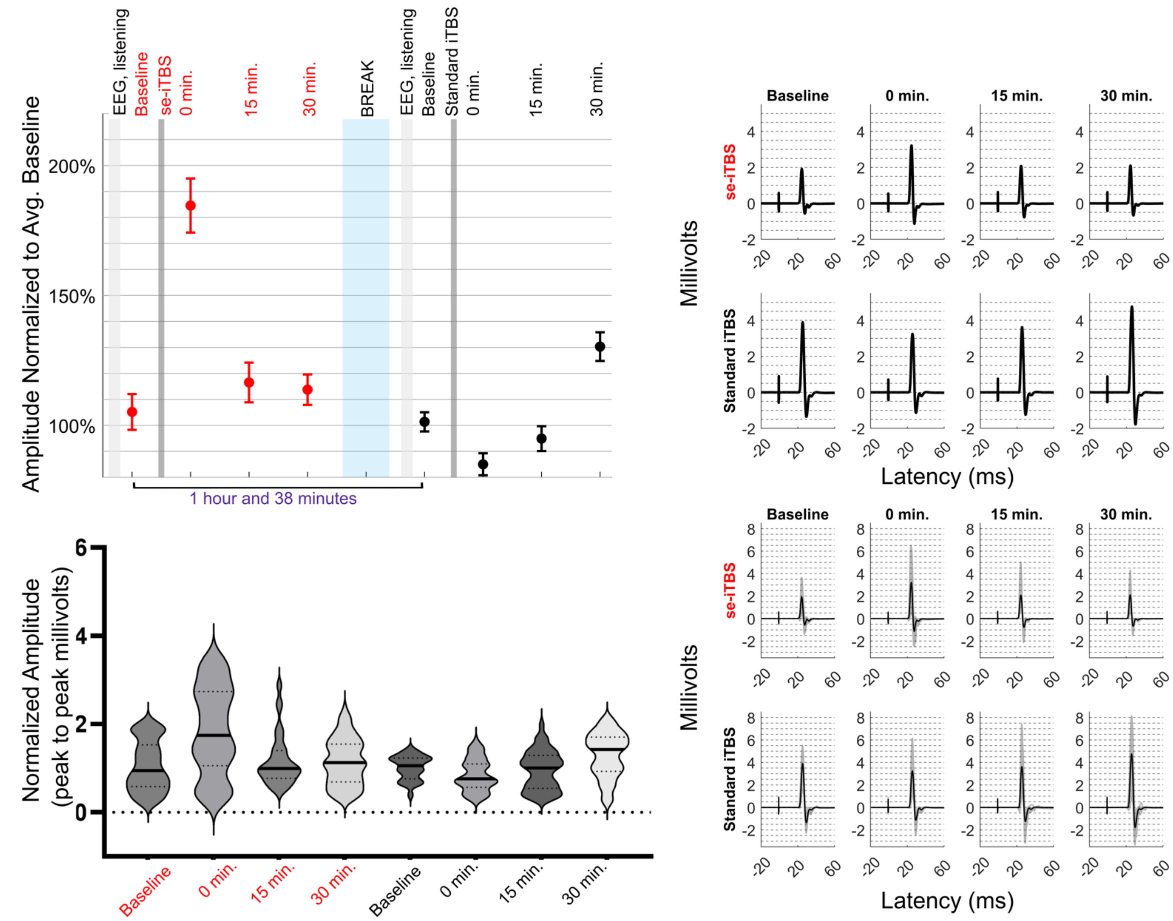


**Figure S11. Participant 7 individual data**

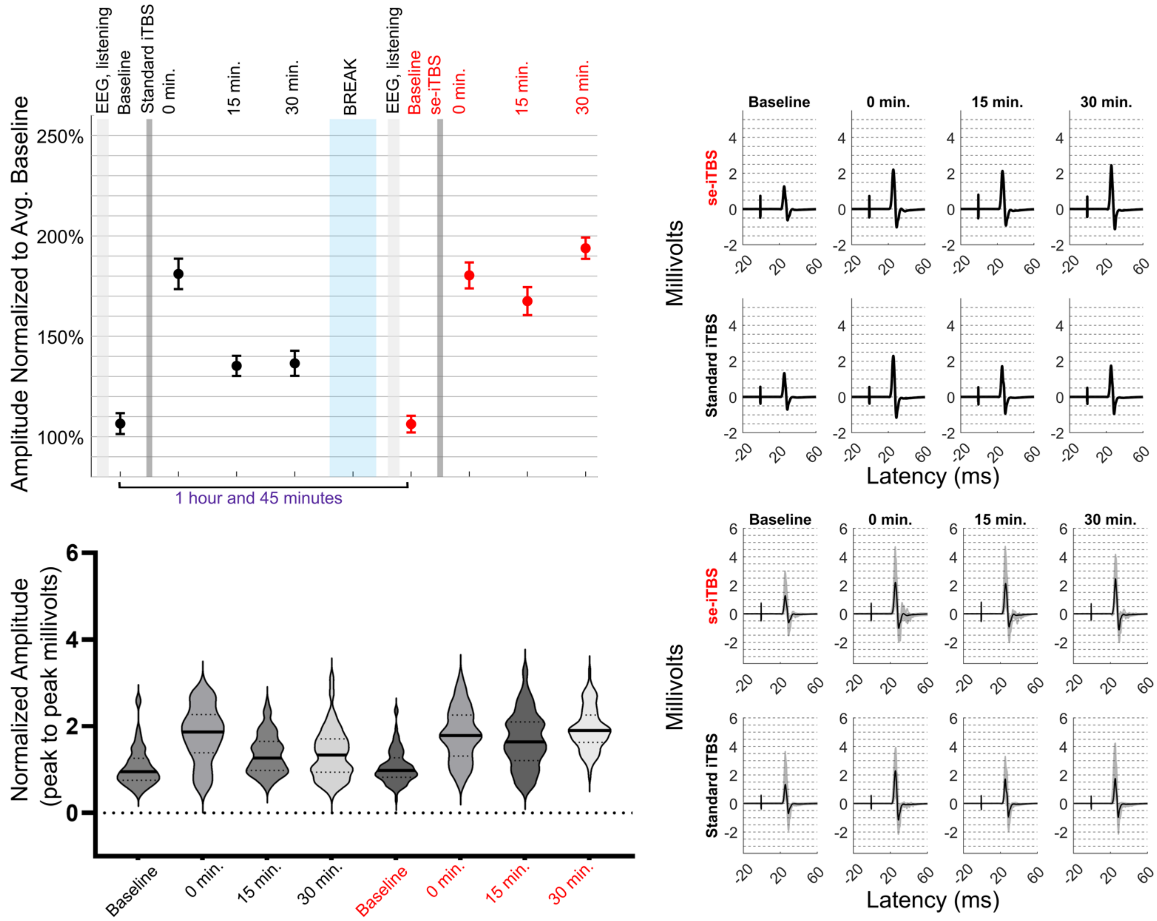


**Figure S12. Participant 8 individual data**

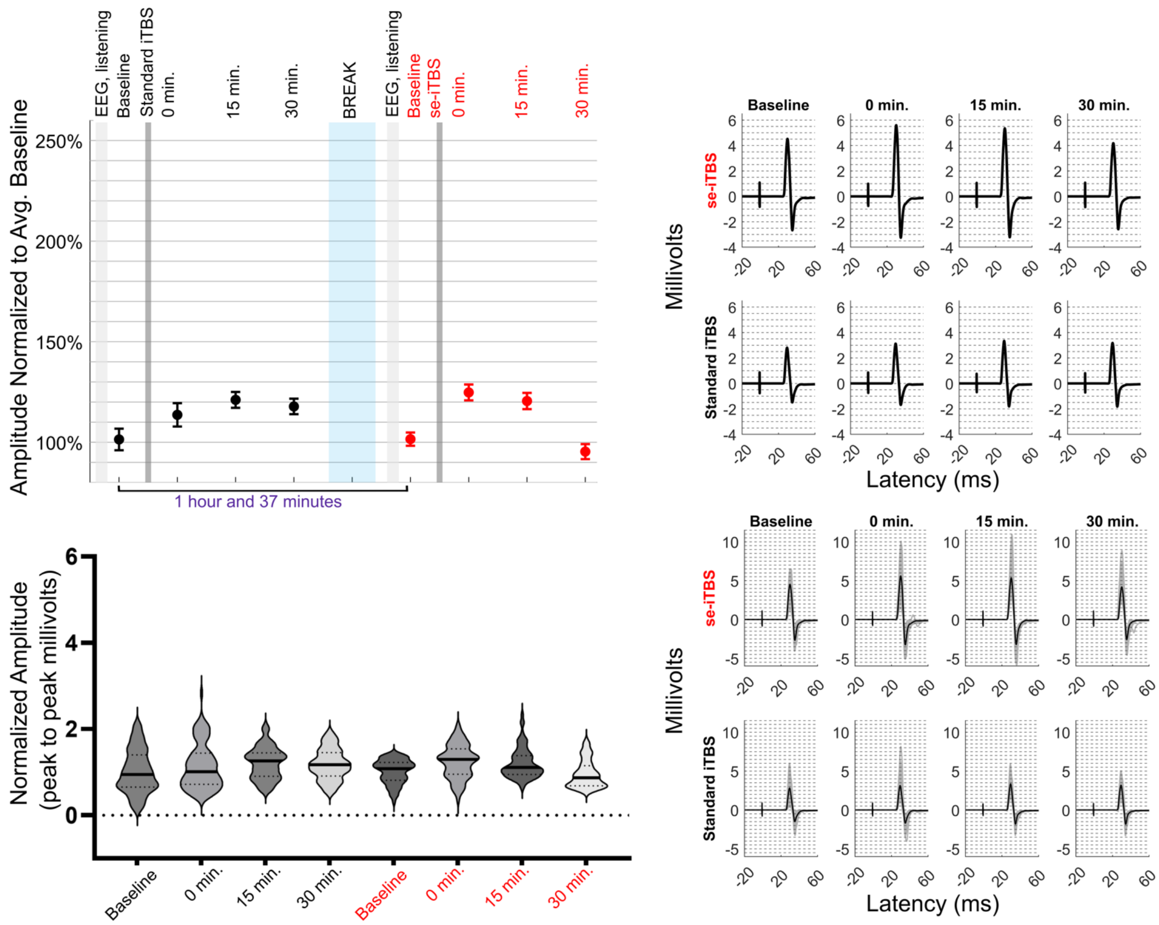


**Figure S13. Participant 9 individual data**

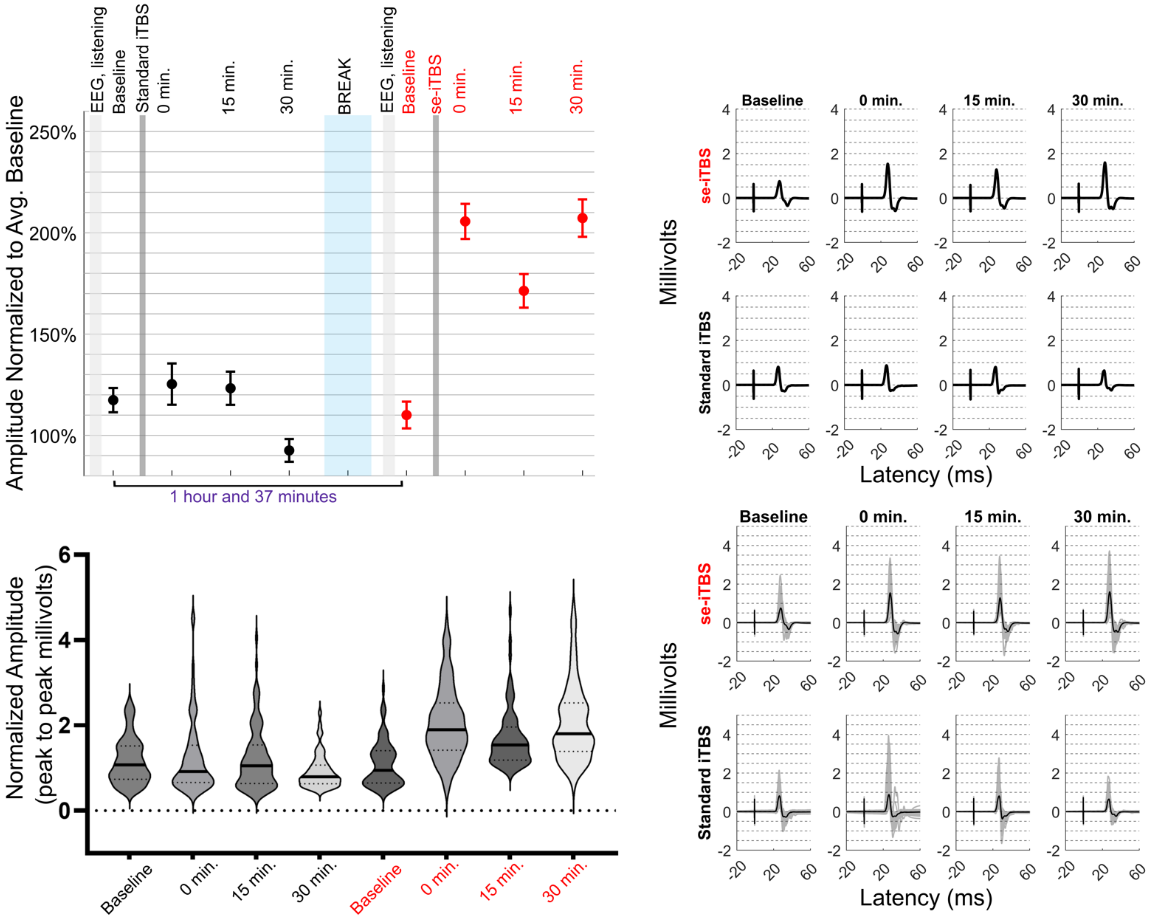


**Figure S14. Participant 10 individual data**

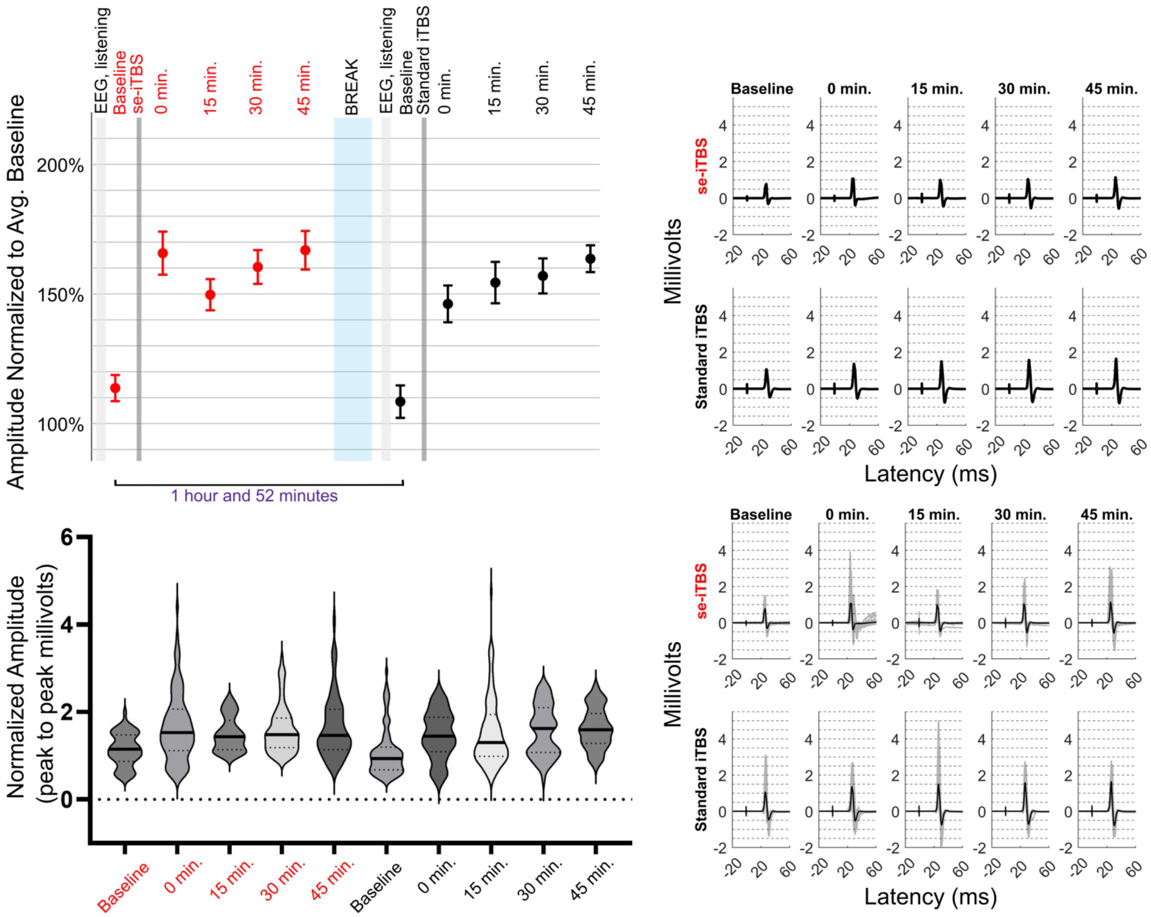


**Figure S15. Participant 11 individual data**

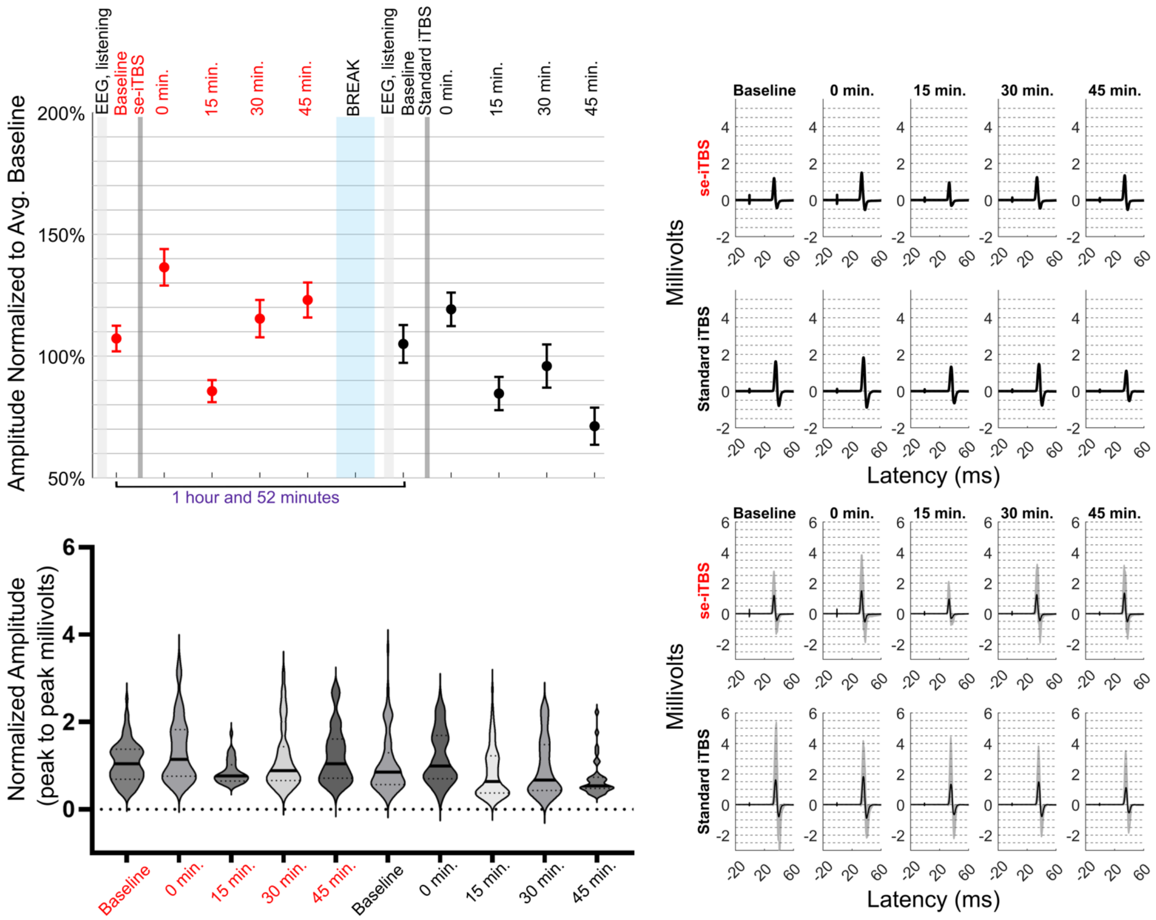


**Figure S16. Participant 12 individual data**

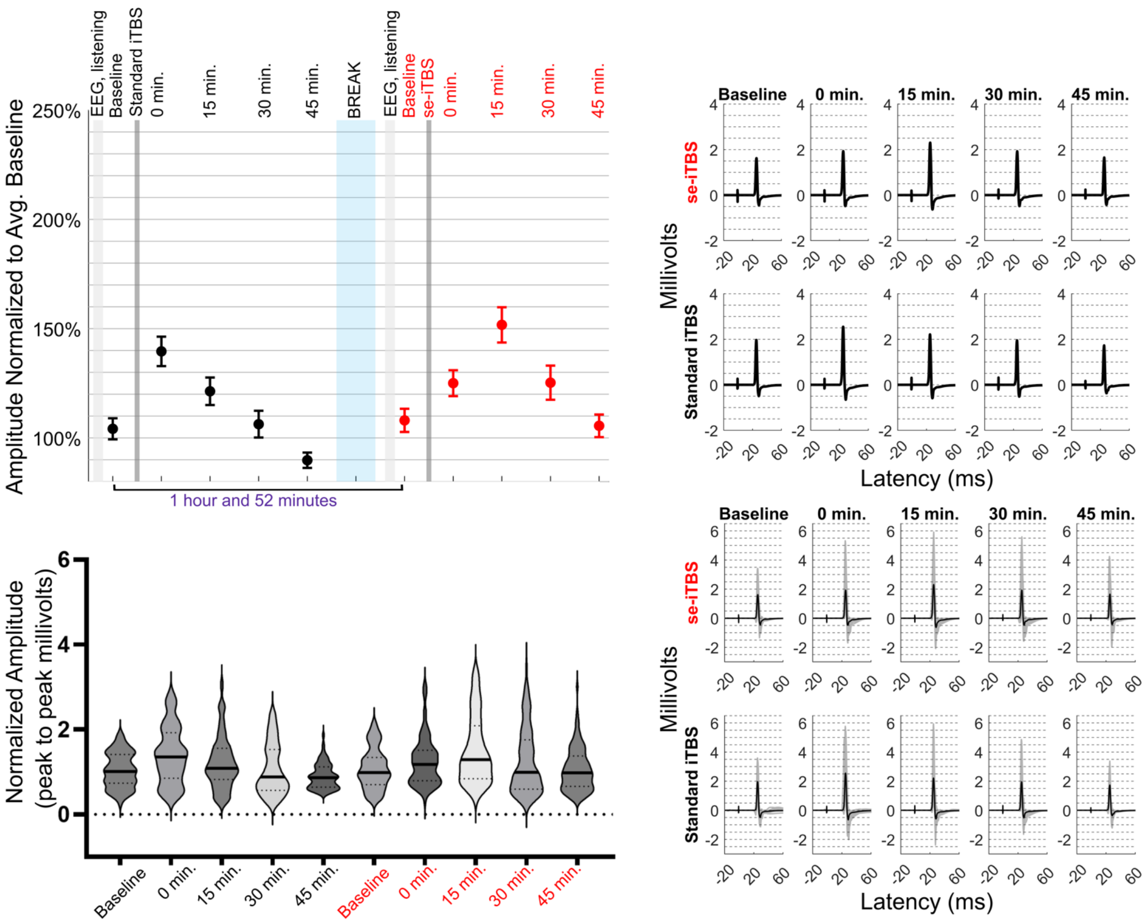


**Figure S17. Participant 13 individual data**

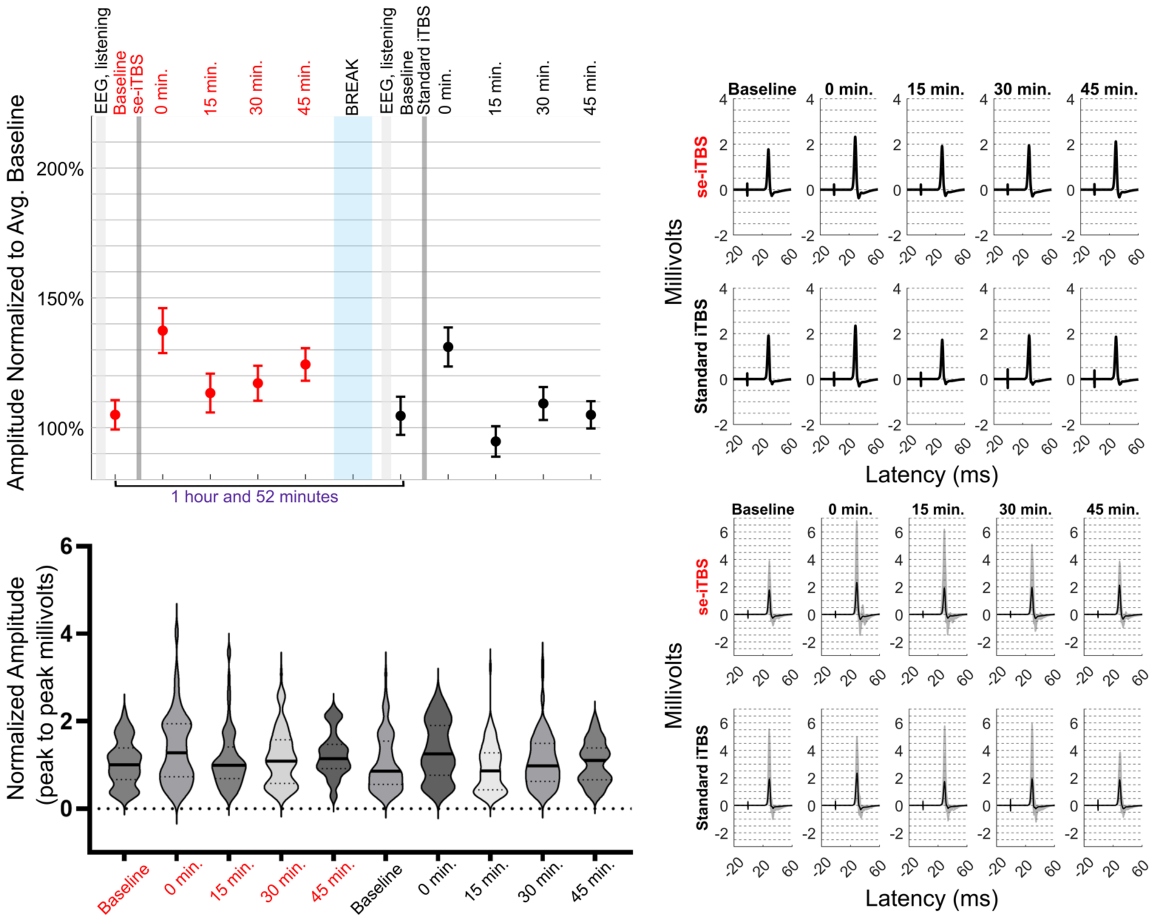


**Figure S18. Participant 14 individual data**

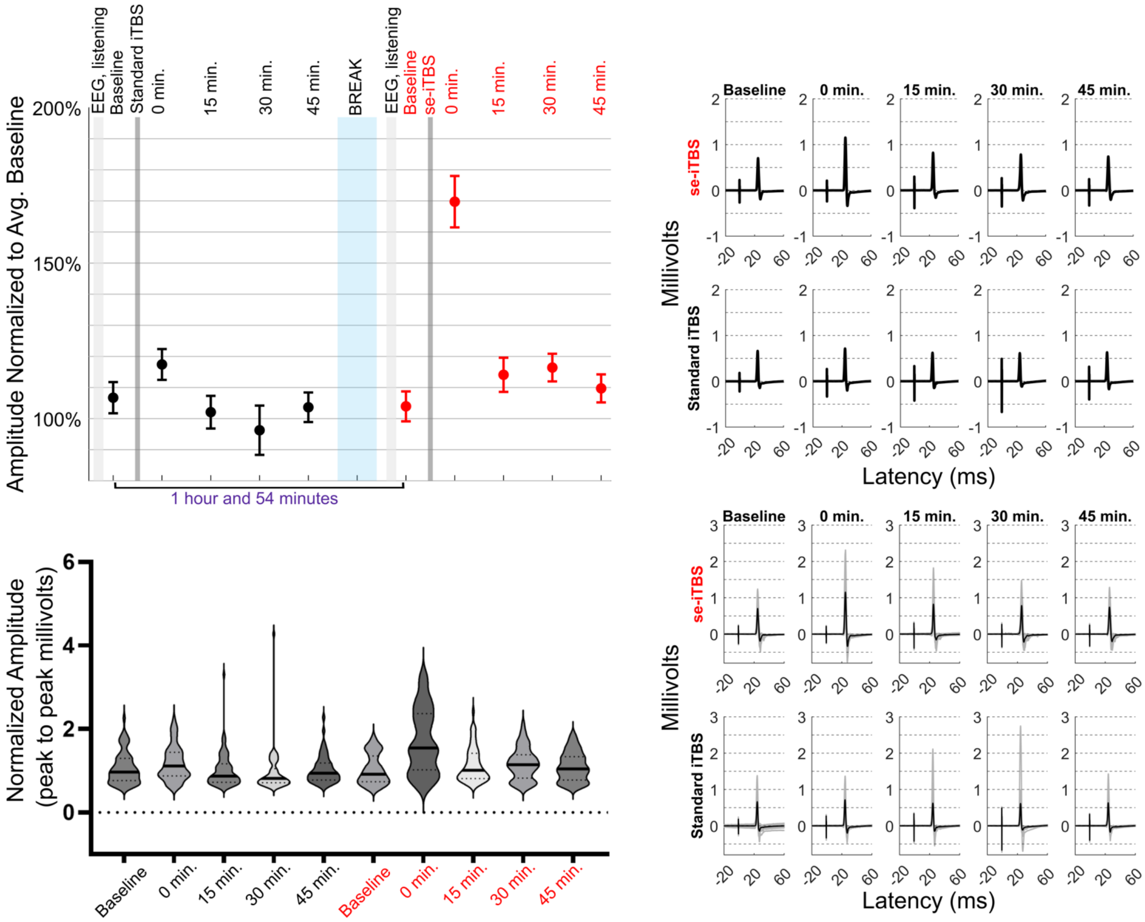


**Figure S19. Participant 15 individual data**

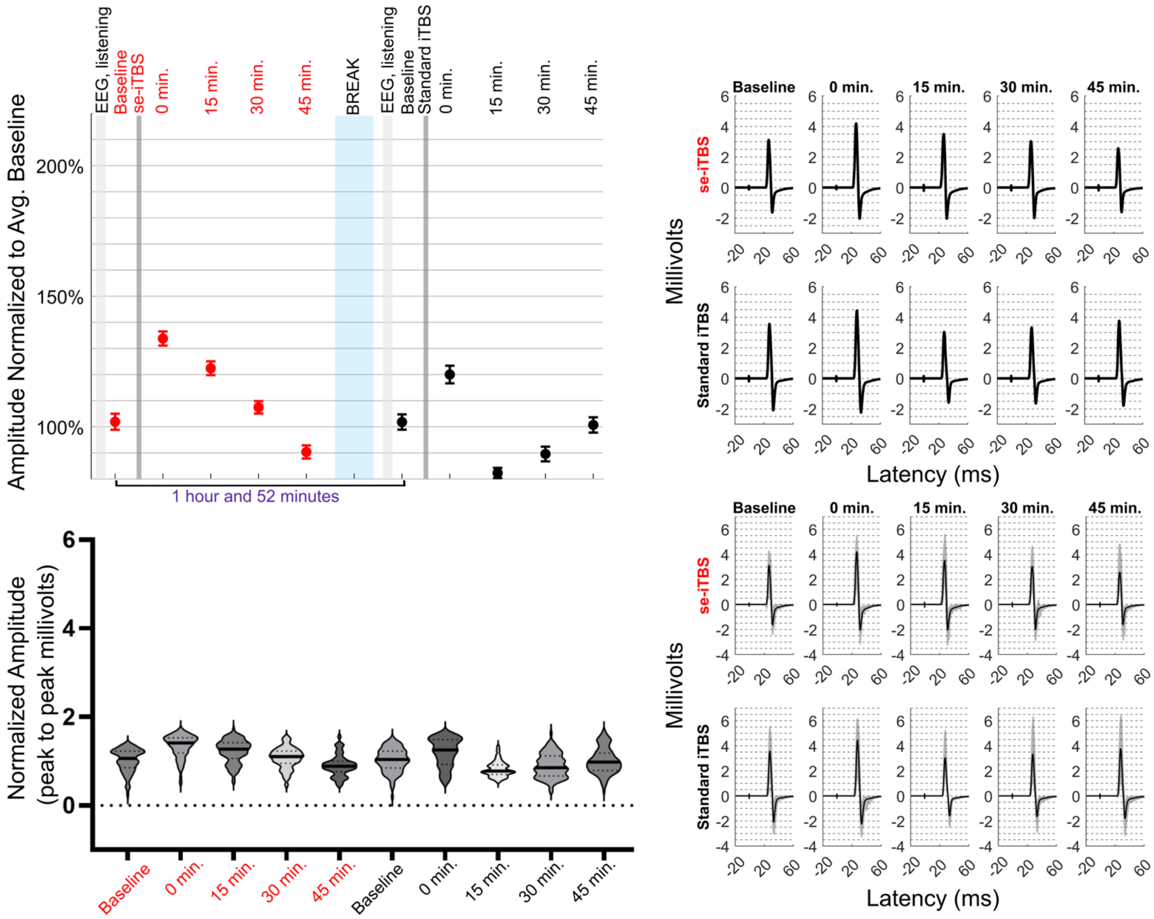


**Figure S20. Participant 16 individual data**

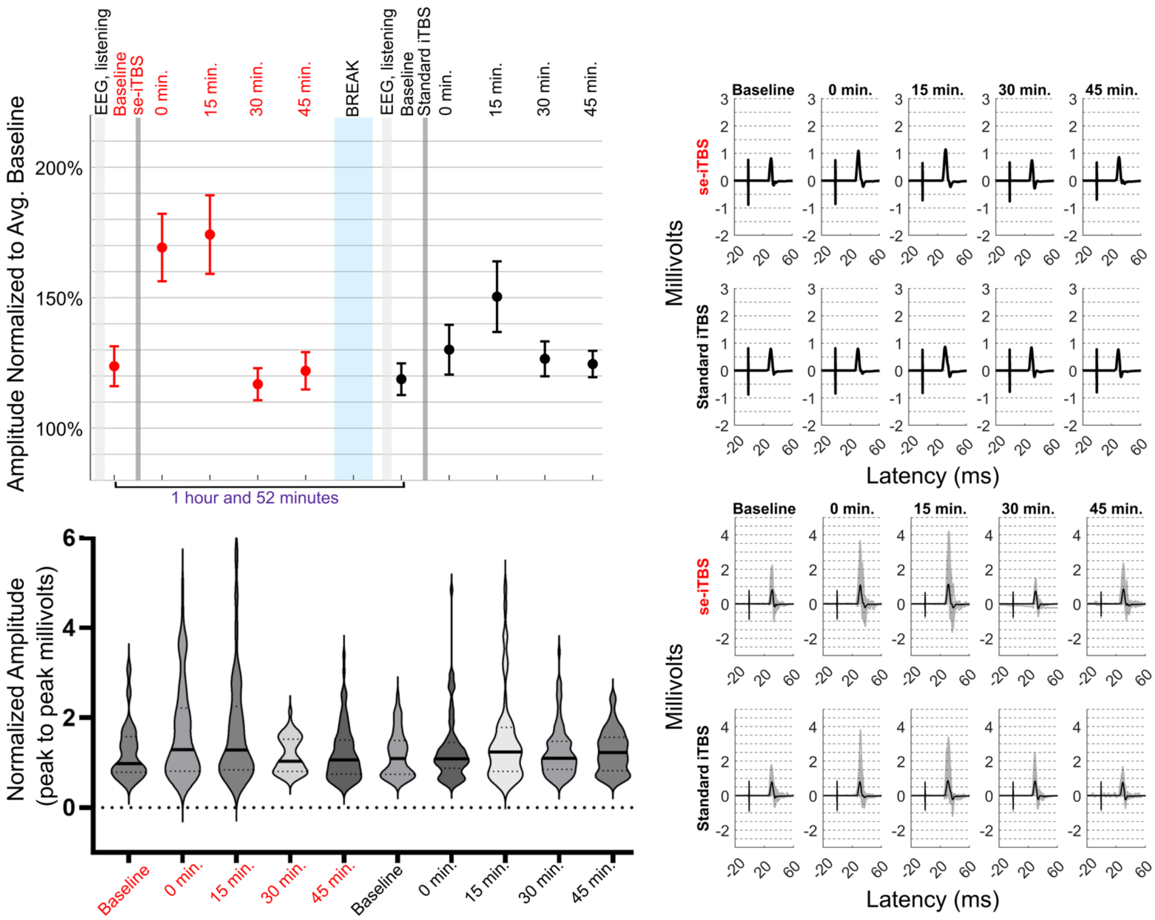


**Figure S21. Participant 17 individual data**

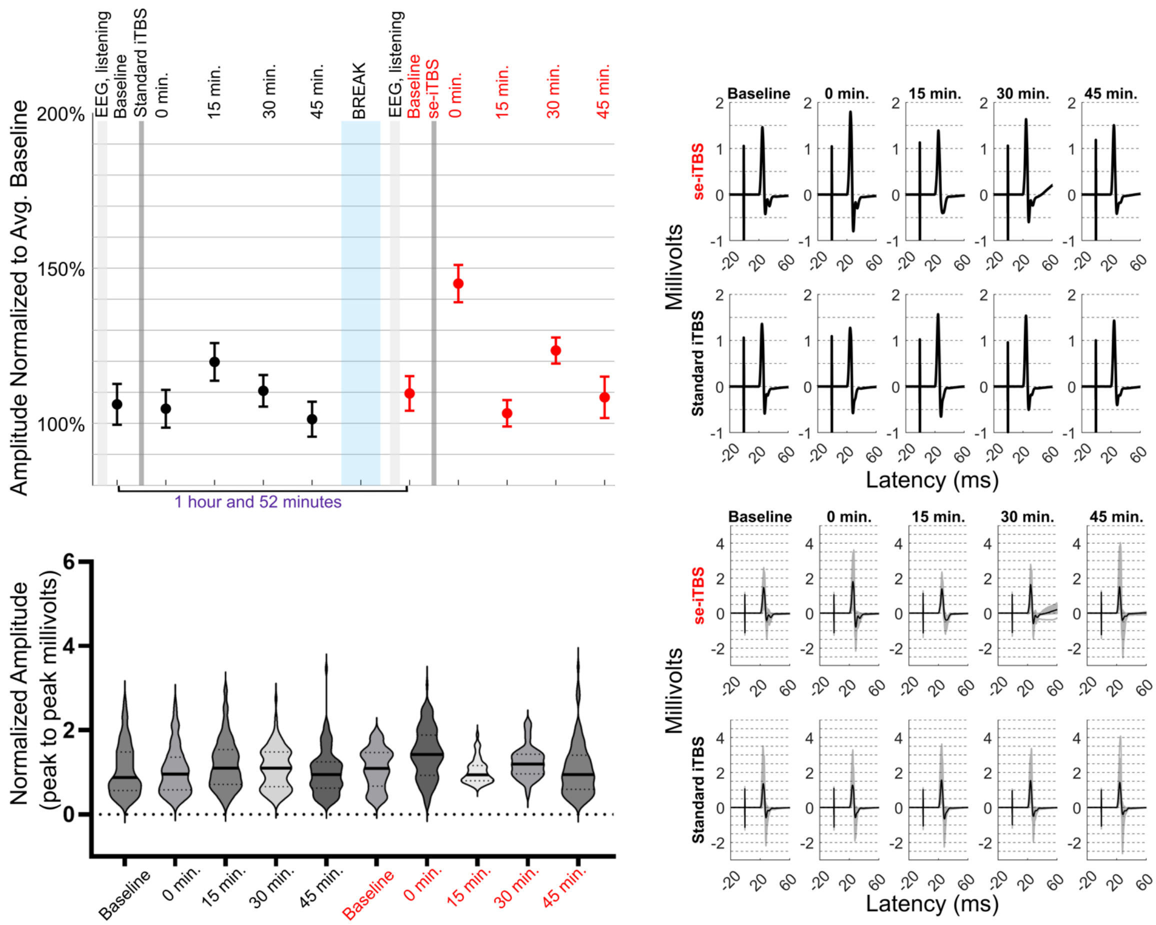


**Figure S22. Participant 18 individual data**

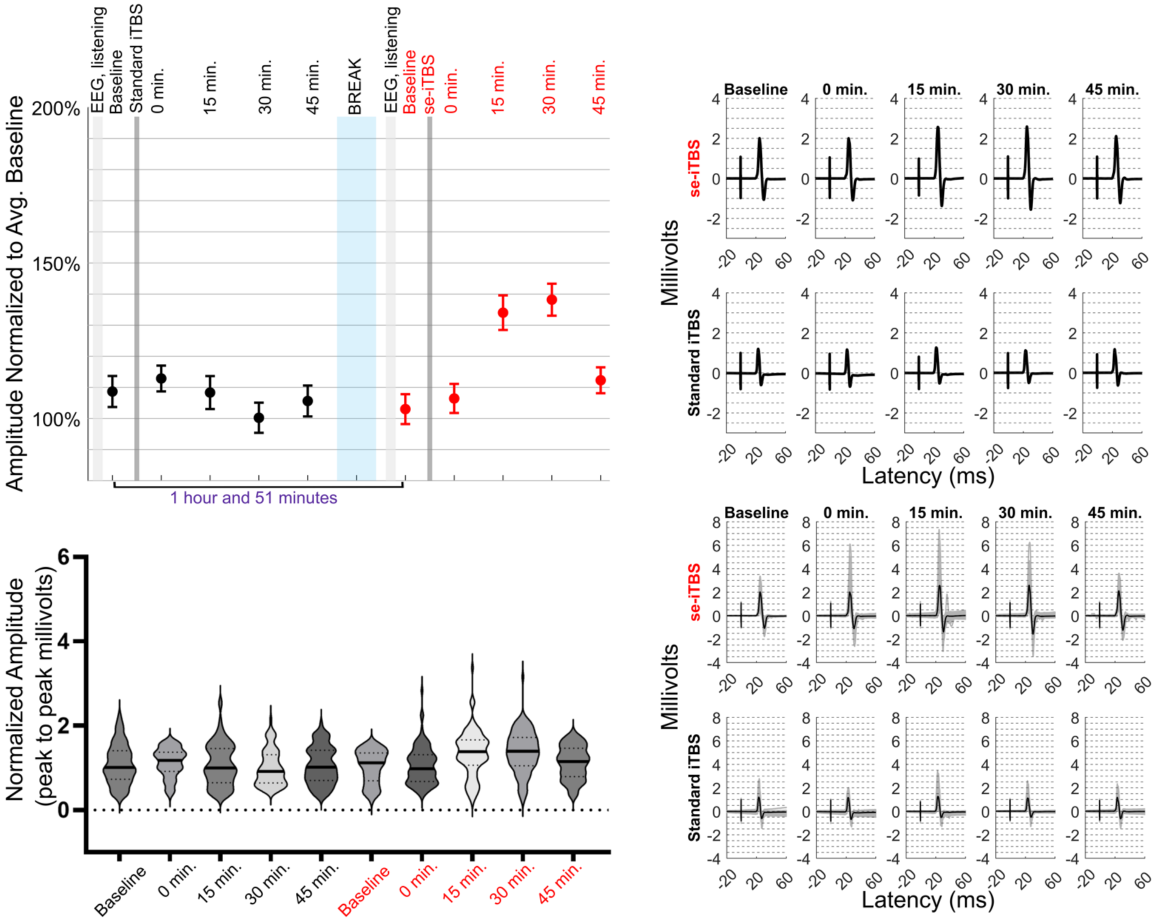


**Figure S23. Participant 19 individual data**

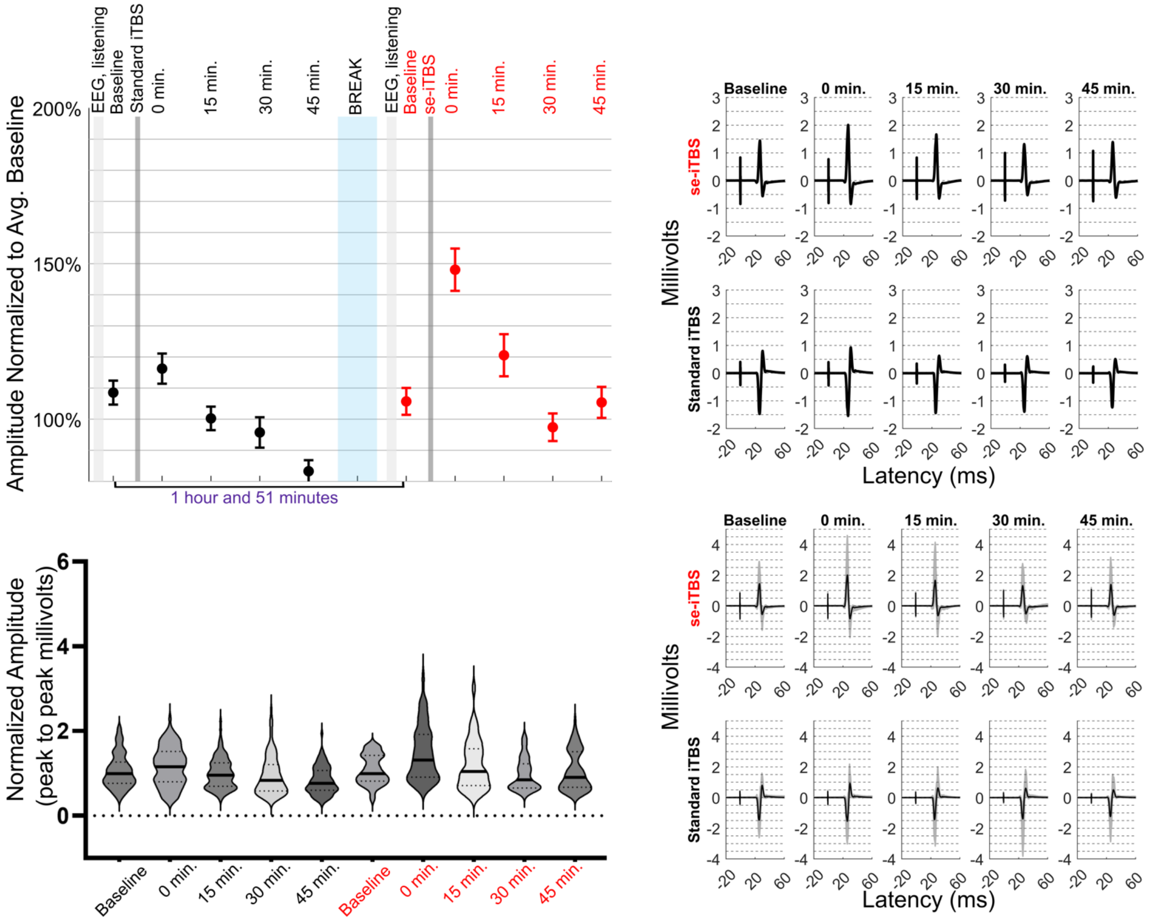


**Figure S24. Participant 20 individual data**
